# A Sac7-Rho1 axis at the plasma membrane controls clathrin-independent endocytosis

**DOI:** 10.64898/2026.07.08.737308

**Authors:** La’Tila Abbott-Wilson, Daniel J. Rioux, Parth R. Patel, Derek C. Prosser

## Abstract

In eukaryotes, our understanding of clathrin-independent endocytosis (CIE) lags far behind that of clathrin-mediated endocytosis (CME). CIE plays key roles in internalizing receptors, viruses, bacterial toxins, and pathogens; thus, deeper mechanistic insights are critical for understanding cellular strategies for plasma membrane regulation. Yeast CIE requires a signal relay between the stress sensor Mid2, the guanine nucleotide exchange factor (GEF) Rom1, the Rho1 GTPase, and the formin Bni1. While GEFs promote GTPase activity, GTPase-activating proteins (GAPs) conversely stimulate nucleotide hydrolysis and GTPase inactivation. Here, we provide new insight into CIE, adding the RhoGAP Sac7 as a regulator. *SAC7* deletion in CME-deficient cells improved cargo internalization, and Sac7 localizes primarily to the mother cortex. Cells lacking *SAC7* accumulate active Rho1 and retain Bni1 at the plasma membrane, where Bni1 retention may subsequently enhance actin assembly needed for CIE. Our results thus demonstrate that Sac7 negatively regulates CIE by restricting cortical Rho1 activity.

## Introduction

Endocytosis, or internalization from the plasma membrane (PM), is critical for regulating cellular communication, PM quality control, and nutrient uptake in eukaryotes. Mechanistically, endocytosis requires cargo concentration, membrane deformation, and scission of a transport carrier. Clathrin-mediated endocytosis (CME) is the best-studied internalization pathway, and involves coordinated action of 40-60 proteins acting in modular waves to regulate endocytic site maturation (Boettner et al., 2012; Kaksonen et al., 2003; Kaksonen et al., 2005; Taylor et al., 2011). Clathrin plays a key role by assembling into a polymeric coat that stabilizes membrane bending, working with adaptors to concentrate cargo at clathrin-coated pits (Kaksonen and Roux, 2018; Reider and Wendland, 2011). The modular nature of CME likely ensures appropriate cargo loading and fidelity of internalization (Kaksonen et al., 2003; Kaksonen et al., 2005).

Separate from CME, numerous clathrin-independent endocytosis (CIE) pathways are known (Mayor et al., 2014; Rioux and Prosser, 2023). Whereas mechanisms including phagocytosis, macropinocytosis, ultrafast endocytosis (UFE) and activity-dependent bulk endocytosis (ADBE) facilitate internalization of larger structures (Kokotos et al., 2018; Watanabe et al., 2013a; Watanabe et al., 2013b), CIE also includes caveolae and pathways involving Rho- and Arf-family GTPases thought to generate smaller vesicles (Rioux and Prosser, 2023; Wang et al., 2009). While CIE pathways can internalize a substantial fraction of the PM, they remain mechanistically obscure, hampered by a scarcity of pathway-specific cargos and, until recently, a lack of genetically tractable model systems. Elucidating mechanisms and physiological functions of CIE pathways, their contributions to cargo internalization, and if/how CME and CIE integrate, is fundamental to our understanding of how cells remodel their PM in response to physiological changes.

Previous studies in the budding yeast *Saccharomyces cerevisiae* identified the first fungal CIE pathway; we now know that CIE is conserved in other yeast and filamentous fungi (Alvaro et al., 2014; Apel et al., 2017; Epp et al., 2013; Martzoukou et al., 2017; Prosser and Wendland, 2012; Prosser et al., 2011; Woodard et al., 2023). Using CME-deficient strains lacking the four clathrin-binding adaptors Ent1, Ent2, Yap1801, and Yap1802, we identified a pathway that relied on the actin-modulating small GTPase Rho1, its guanine nucleotide exchange factor (GEF) Rom1, and its effector, the formin Bni1 (Maldonado-Báez et al., 2008; Prosser et al., 2011). The Rho1-dependent pathway promotes internalization of multiple cargos and acts in virtually all CME-deficient mutants tested, including loss-of-function of actin stabilizers (*sac6*Δ *scp1*Δ), actin nucleators (*las17*Δ), scaffolding proteins (*end3*Δ), and clathrin (*chc1*Δ) (Alvaro et al., 2014; Apel et al., 2017; Prosser et al., 2011). In *S. cerevisiae* CIE, the formin Bni1 elongates unbranched actin filaments, which are mutually exclusive from Arp2/3-generated branched networks at cortical actin patches that mark CME sites (Burke et al., 2014; Evangelista et al., 2001; Pruyne et al., 2002), demonstrating that the Rho1-dependent pathway is distinct from CME. Subsequent studies expanded the set of yeast CIE proteins to include cargo-selective α-arrestins and the Fes/CIP4 homology-Bin/amphiphysin/Rvs (F-BAR) protein Syp1 (also involved in CME), the type V myosin Myo2, microtubules, dynein/dynactin, and cortical microtubule transport and capture machinery (Alvaro et al., 2014; Apel et al., 2017; Reider et al., 2009; Woodard et al., 2023). Most CIE-involved proteins are conserved, suggesting elements of the yeast Rho1 pathway contribute to endocytosis in other cell types; indeed, the Rho1 homolog RhoA promotes compensatory CIE in bladder umbrella cells and uptake of the IL-2 receptor (Khandelwal et al., 2010; Lamaze et al., 2001). Similarly, formins contribute to neuronal UFE, while microtubules are involved in fast endophilin-mediated endocytosis (FEME) (Casamento and Boucrot, 2020; Soykan et al., 2017; Tyckaert et al., 2022). Our current understanding of CIE suggests that many pathways depend on sets of proteins that partially overlap between different CIE modalities, or with CME, and may thus share key mechanistic similarities (Rioux and Prosser, 2023).

Localized recruitment and/or activity is critical for small GTPase function. Two enzyme classes primarily regulate small GTPases: GEFs and GTPase-activating proteins (GAPs) (Mizuno-Yamasaki et al., 2012). GEFs promote activation via exchange of GDP for GTP, leading to association with effectors that perform downstream functions. In contrast, GAPs stimulate GTP hydrolysis, resulting in inactivation of the GTPase. These opposing roles enable activation and inactivation cycles, where GEF and GAP localization can contribute to spatial and temporal GTPase regulation through incompletely understood mechanisms. We demonstrated that high-copy expression of the Rho1 GEF *ROM1* promoted CIE in yeast (Prosser et al., 2011), and so we asked whether GAPs negatively regulate Rho1 at the PM to control CIE. We show here that loss of the GAP Sac7, but not other Rho1 GAPs, restores cargo internalization in CME-deficient cells by promoting CIE. Unexpectedly, Sac7 localization at the mother cortex is necessary for polarization of active Rho1 to the bud tip, as *sac7*Δ causes aberrant accumulation of Rho1-GTP at the mother PM. In turn, increased active Rho1 at the PM stabilizes cortical Bni1 and promotes additional assembly of small, actin cable-like structures. Our results demonstrate that Sac7 negatively regulates Rho1 function in CIE by stimulating localized GTPase inactivation. We posit that this GAP-dependent regulation may contribute to establishment and/or maintenance of appropriate Rho GTPase activity gradients.

## Results

### Loss of the Rho1 GAP SAC7 restores endocytosis in CME-deficient cells

Previous results demonstrated that high-copy *RHO1* and its GEF, *ROM1*, promote CIE in CME-deficient yeast strains (Prosser et al., 2011). Since GEF upregulation increases Rho1 activity, we reasoned that deletion of GAPs, which stimulate GTP hydrolysis, might also promote CIE by preventing PM inactivation of Rho1. *S. cerevisiae* possesses four Rho1 GAPs (Bag7, Bem2, Lrg1, and Sac7; hereafter referred to as RhoGAPs), which differentially contribute to Rho1-dependent functions (Bickle et al., 1998; Lorberg et al., 2001; Schmidt et al., 2002; Watanabe et al., 2001). Bem2, Lrg1 and Sac7 regulate cell wall integrity (CWI) downstream of stress sensors including Wsc1 and Mid2, as well as through Tor2 signaling (Lorberg et al., 2001; Schmidt et al., 2002; Watanabe et al., 2001). The CWI pathway activates protein kinase C (Pkc1), the MAP kinase Slt2, and the formin Bni1 to polarize actin cables and deliver repair enzymes to damage sites (Helliwell et al., 1998; Richthammer et al., 2012). Separate from CWI, Bag7 and Sac7 promote actin reorganization downstream of stress sensors through poorly-understood mechanisms. Together, Lrg1 and Sac7 are the major RhoGAPs, while Bag7 and Bem2 likely play minor or specialized roles (Schmidt et al., 2002; Yoshida et al., 2009).

To assess contributions to CME and CIE, we examined deletions of each RhoGAP alone, and in cells lacking the clathrin-binding endocytic adaptors Ent1, Ent2, Yap1801, and Yap1802 (4Δ). Yeast CME is efficient in cells expressing any single adaptor (Maldonado-Báez et al., 2008); thus, we used both WT and 4Δ+Ent1 backgrounds expressing plasmid-borne *ENT1* (hereafter referred to as 4Δ^ON^) as controls with functional CME. In contrast, 4Δ+ENTH1 cells (hereafter referred to as 4Δ^OFF^), which express the phosphatidylinositol 4,5-bisphosphate [PI(4,5)P_2_]-binding epsin N-terminal homology (ENTH) domain of Ent1 for viability due to synthetic lethality of *ent1*Δ *ent2*Δ, are deficient in CME and show PM retention of endocytic cargos (Aguilar et al., 2006; Maldonado-Báez et al., 2008). Additionally, 4Δ^OFF^ cells are temperature-sensitive, displaying reduced growth at 35.5°C and inviability at 37°C (Maldonado-Báez et al., 2008; Prosser et al., 2011).

First, we examined effects of RhoGAP deletion on temperature-dependent growth in 4Δ^OFF^. As expected, 4Δ^OFF^ cells showed reduced growth at 35.5°C and above compared to WT and 4Δ^ON^ (Fig. 1A). Deletion of *BAG7*, *BEM2*, and *LRG1* did not alter temperature sensitivity in CME-deficient 4Δ^OFF^ cells, which showed similar growth patterns to 4Δ^OFF^ controls without RhoGAP deletion. As observed previously, *bem2*Δ alone resulted in temperature sensitivity (Bender and Pringle, 1991), even though *bem2*Δ 4Δ^ON^ cells showed improved growth compared to *bem2*Δ. In contrast to the other GAPs, *sac7*Δ restored growth at high temperatures in 4Δ^OFF^ to levels resembling *sac7*Δ and *sac7*Δ 4Δ^ON^ controls, as well as to WT and 4Δ^ON^ controls. These data suggest that loss of Sac7, but not other GAPs, improves temperature-dependent growth of 4Δ^OFF^ cells.

**Figure 1:**
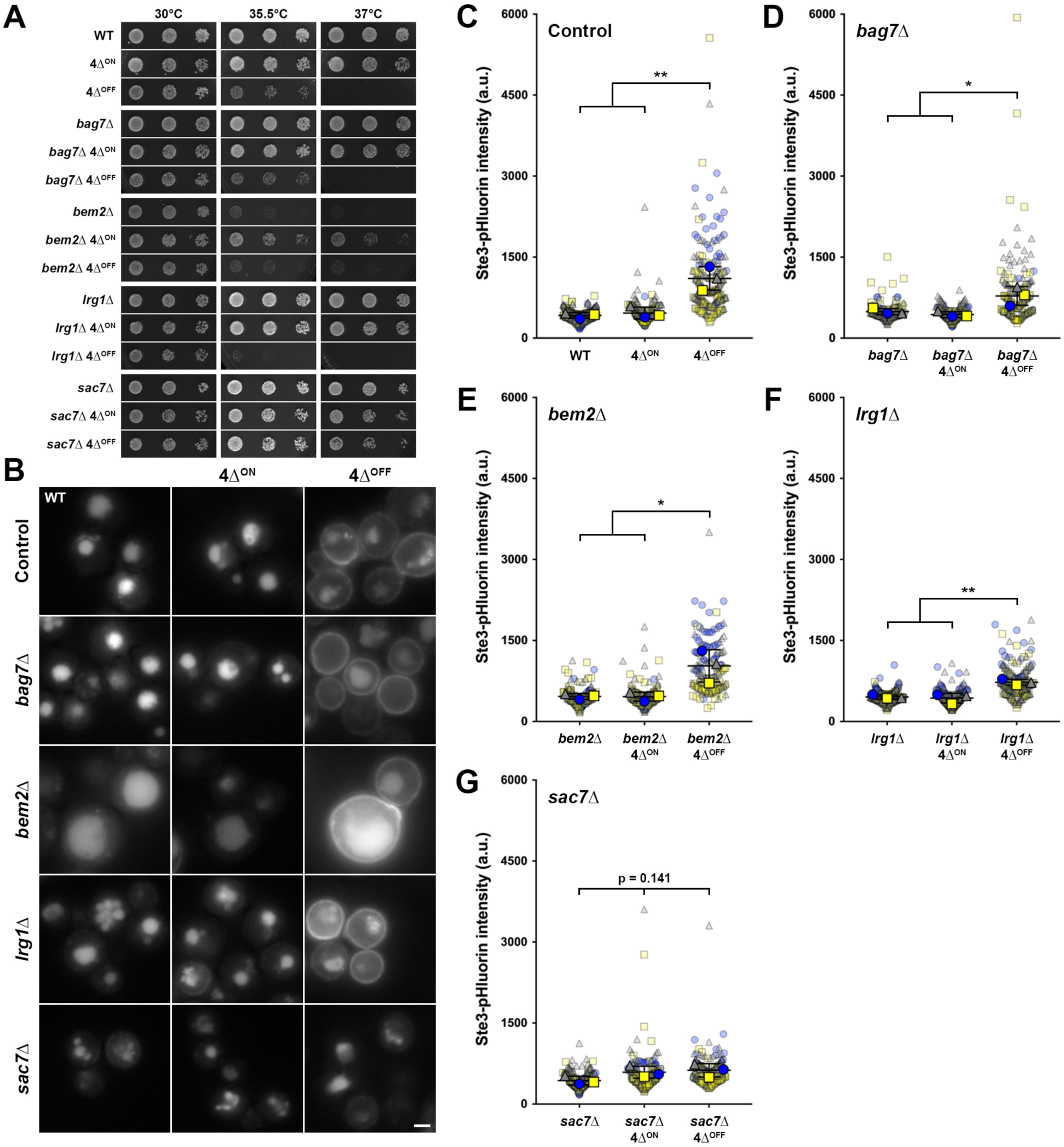
Effect of Rho1 GAP deletion on temperature-dependent growth and endocytosis in adaptor-deficient cells. (A) WT (control), 4Δ^ON^ and 4Δ^OFF^ cells without and with Rho1 GAP deletions as indicated were grown to mid-logarithmic phase, and identical five-fold serial dilution series were plated on YPD medium for 3-4 d at 30, 35.5, and 37°C. (B) Fluorescence microscopy of cells as described in panel *A*, expressing Ste3-GFP. Scale bar, 2 µm. (C-G) Quantification of endocytic capacity using Ste3-pHluorin. Whole-cell fluorescence intensity of WT, 4Δ^ON^, and 4Δ^OFF^ cells in control (no Rho1 GAP deletion; C), *bag7*Δ (D), *bem2*Δ (E), *lrg1*Δ (F), and *sac7*Δ (G) strains was measured for n=3 biological replicates, with 36-105 cells measured per condition for each trial. One-way ANOVA with Tukey’s multiple comparisons test, * p<0.05; ** p<0.01.

We next examined steady-state localization of the pheromone receptor Ste3 as a representative cargo that undergoes constitutive endocytosis and vacuolar targeting in WT and 4Δ^ON^. We previously observed Ste3-GFP PM retention in 4Δ^OFF^ cells and internalization via CIE in 4Δ^OFF^ cells expressing high-copy *ROM1*, as seen for numerous other cargos (Apel et al., 2017; Prosser et al., 2011; Prosser et al., 2015). In WT, *bag7*Δ, *bem2*Δ, *lrg1*Δ, and *sac7*Δ, as well as 4Δ^ON^ and RhoGAP-deficient 4Δ^ON^ cells, Ste3-GFP localized prominently to the vacuole and was barely detectable at the PM, indicating that GAP deletions did not impair Ste3 internalization when CME is functional (Fig. 1B). Loss of *BEM2* resulted in abnormally large and fragile-appearing cells, consistent with previous studies (Marquitz et al., 2002). As expected, Ste3-GFP showed PM retention in 4Δ^OFF^, consistent with defective endocytosis. For *bag7*Δ, *bem2*Δ, and *lrg1*Δ in the 4Δ^OFF^ background, Ste3-GFP was PM-retained; however, *sac7*Δ 4Δ^OFF^ cells had improved Ste3-GFP internalization and vacuolar delivery, indicating that loss of Sac7 restores endocytosis in 4Δ^OFF^ (Fig. 1B).

To quantify relative endocytic capacity, we used the same strains, but expressing Ste3 with a C-terminal pHluorin tag. GFP fluorescence persists after cargo delivery to the vacuole; thus, whole-cell Ste3-GFP fluorescence intensity does not reliably report differences in cargo internalization. In contrast, pHluorin becomes quenched in acidic environments such as the vacuole lumen, and fluorescence of cargos bearing a cytoplasmic pHluorin tag is only visible at the PM and occasionally on endosomes (Miesenböck et al., 1998; Prosser et al., 2010; Prosser et al., 2016). Thus, WT and 4Δ^ON^ cells with functional CME have low Ste3-pHluorin intensity, while 4Δ^OFF^ are comparatively bright (Fig. 1C). Elevated fluorescence in 4Δ^OFF^ compared to WT and 4Δ^ON^ backgrounds was also observed in *bag7*Δ, *bem2*Δ, and *lrg1*Δ strains, confirming that deletion of these GAPs did not improve endocytosis in CME-deficient cells (Fig. 1D-F). Consistent with improved Ste3-GFP vacuolar delivery, *sac7*Δ 4Δ^OFF^ had Ste3-pHluorin intensity similar to *sac7*Δ and *sac7*Δ 4Δ^ON^, indicating that *sac7*Δ improves endocytosis in CME-deficient cells (Fig. 1G).

### Sac7 operates in the Rho1-dependent CIE pathway

Since Sac7 is a RhoGAP, Ste3-GFP internalization in *sac7*Δ 4Δ^OFF^ cells is likely through Rho1-dependent CIE, but could also occur by reactivating CME via an unknown mechanism. Previous results showed that Bni1 is required downstream of Rho1 for CIE (Prosser et al., 2011). We reasoned that if Sac7 acts in the same pathway as Rho1 and Bni1, additional deletion of *BNI1* would result in an epistatic interaction to block the rescue of temperature sensitivity and Ste3 internalization in *sac7*Δ 4Δ^OFF^. As before, *sac7*Δ 4Δ^OFF^ grew at elevated temperature similar to *sac7*Δ and *sac7*Δ 4Δ^ON^ (Figs. 1A and 2A), while *bni1*Δ 4Δ^OFF^ cells defective in both CME and CIE were inviable at 35.5°C and above (Fig. 2A). Both *bni1*Δ and *bni1*Δ 4Δ^ON^ grew at all temperatures examined. Consistent with the predicted epistasis, *bni1*Δ *sac7*Δ 4Δ^OFF^ cells were inviable at 35.5°C and above, unlike the respective control and 4Δ^ON^ conditions.

**Figure 2:**
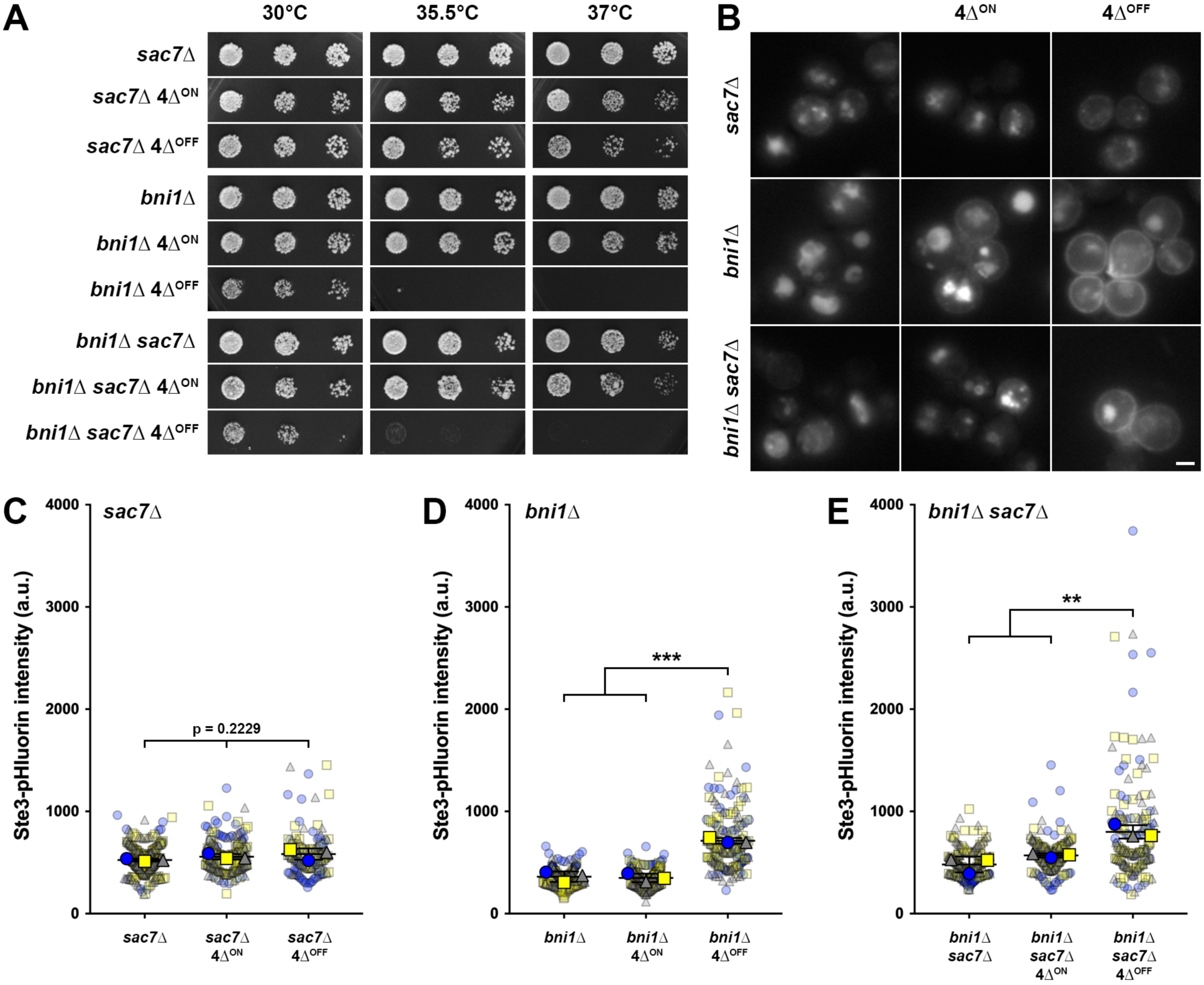
Assessment of the role of Sac7 in clathrin-independent endocytosis. (A) Control, 4Δ^ON^ and 4Δ^OFF^ cells in *sac7*Δ, *bni1*Δ, and *sac7*Δ *bni1*Δ backgrounds were grown to mid-logarithmic phase, and identical five-fold serial dilution series were plated on YPD medium for 3-4 d at 30, 35.5, and 37°C. (B) Fluorescence microscopy of cells as described in panel *A*, expressing Ste3-GFP. Scale bar, 2 µm. (C-E) Quantification of endocytic capacity of cells as in panel *B*, but expressing Ste3-pHluorin. Whole-cell fluorescence intensity of control, 4Δ^ON^ and 4Δ^OFF^ cells in *sac7*Δ (C), *bni1*Δ (D), or *sac7*Δ *bni1*Δ (E) was measured for n=3 biological replicates, with 41-111 cells measured per condition for each trial. One-way ANOVA with Tukey’s multiple comparisons test, ** p<0.01; *** p<0.001.

In addition to temperature-dependent growth, we compared Ste3 internalization in *sac7*Δ, *bni1*Δ, and *bni1*Δ *sac7*Δ strains. In all WT and 4Δ^ON^ backgrounds, Ste3-GFP internalized efficiently and was targeted to the vacuole (Fig. 2B). As expected, Ste3-GFP was primarily vacuolar in *sac7*Δ 4Δ^OFF^ cells, and showed PM retention in *bni1*Δ 4Δ^OFF^ cells. Consistent with Sac7 acting with Rho1 and Bni1, *bni1*Δ *sac7*Δ 4Δ^OFF^ cells retained Ste3-GFP at the PM, indicating that *sac7*Δ fails to restore endocytosis when the formin that drives CIE is absent. Quantification of Ste3-pHluorin intensity agreed with the qualitative localization data for Ste3-GFP. Whereas Ste3-pHluorin intensity in *sac7*Δ 4Δ^OFF^ was indistinguishable from *sac7*Δ and *sac7*Δ 4Δ^ON^, indicative of restored endocytosis (Fig. 2C), *bni1*Δ 4Δ^OFF^ and *bni1*Δ *sac7*Δ 4Δ^OFF^ cells were significantly brighter than their respective controls (WT and 4Δ^ON^ backgrounds; Fig. 2D-E). Epistasis between Bni1 and Sac7 thus suggests that both proteins act in the same CIE pathway.

### Sac7 GAP activity is required for its regulation of CIE

To demonstrate that Sac7 enzymatic activity is required for CIE, rather than recruitment of other endocytic accessory proteins, we used low-copy plasmids to reintroduce *SAC7^WT^* or mutant *sac7^R173K^*, which lacks the catalytic arginine finger (Ho et al., 2008). We predicted that Sac7^WT^, but not inactive Sac7^R173K^, would effectively undo rescue of temperature sensitivity and endocytosis in *sac7*Δ 4Δ^OFF^. Indeed, *sac7*Δ, *sac7*Δ 4Δ^ON^ and *sac7*Δ 4Δ^OFF^ cells transformed with empty vector were temperature-resistant, while *sac7*Δ 4Δ^OFF^ cells transformed with *SAC7^WT^* grew at 30°C but not at higher temperatures (Fig. 3A), consistent with the model that restoration of Sac7 function impedes CIE activation. In contrast, *sac7*Δ 4Δ^OFF^ expressing *sac7^R173K^* remained temperature-resistant, and in fact grew more robustly than with empty vector. Moreover, *sac7*Δ, *sac7*Δ 4Δ^ON^ and *sac7*Δ 4Δ^OFF^ transformed with empty vector all showed similar vacuolar delivery of Ste3-GFP and low Ste3-pHluorin fluorescence (Fig. 3B-C). Whereas *SAC7^WT^* reduced Ste3-GFP internalization and increased Ste3-pHluorin intensity in *sac7*Δ 4Δ^OFF^, *sac7^R173K^* had no observable effect on cargo internalization compared to empty vector. These data indicate that Sac7 catalytic activity is required for regulation of CIE.

**Figure 3:**
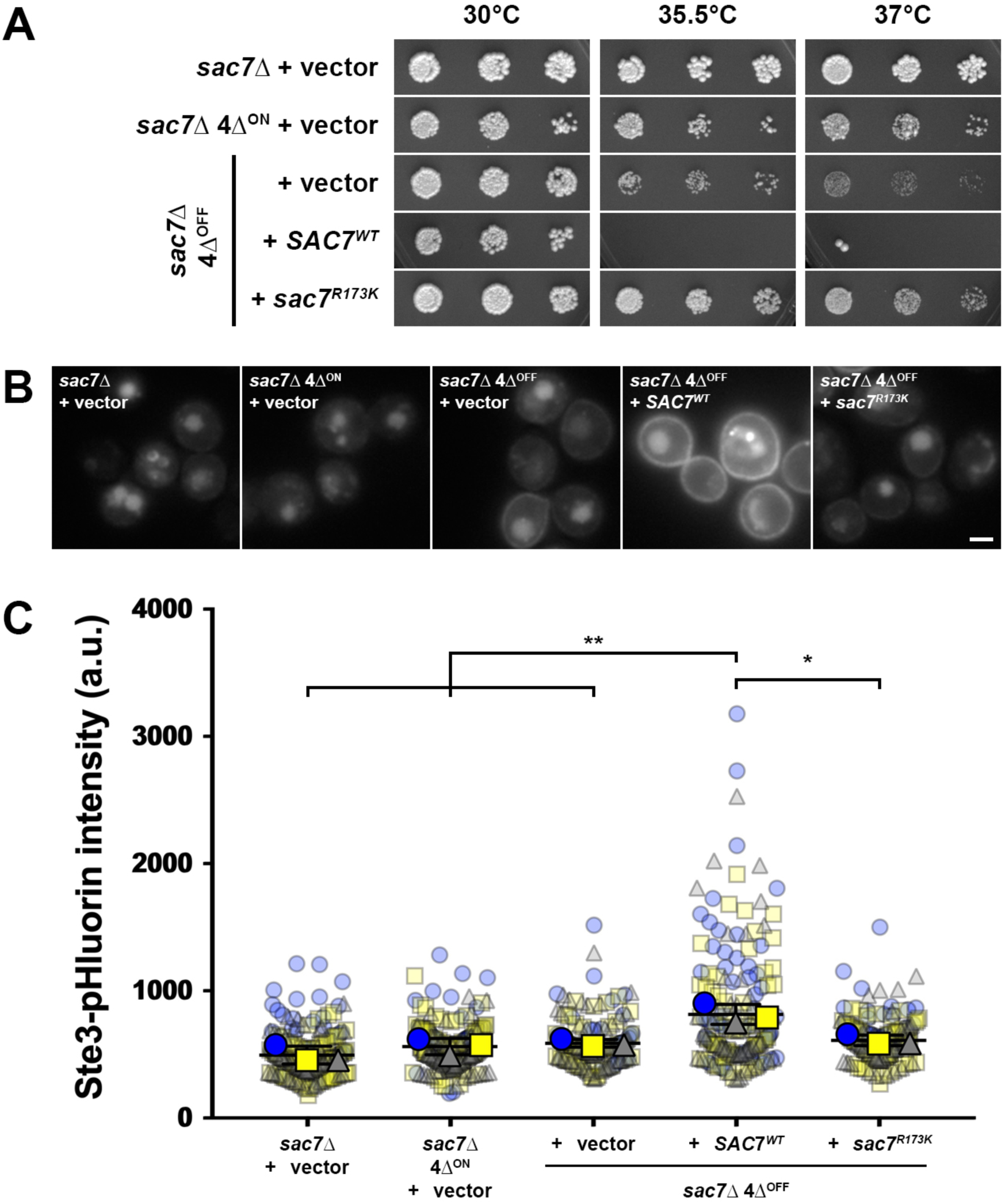
Requirement of Sac7 GAP activity for its role in regulating clathrin-independent endocytosis. (A) *sac7*Δ, *sac7*Δ 4Δ^ON^, and *sac7*Δ 4Δ^OFF^ cells transformed with empty vector or low-copy *SAC7^WT^*or *sac7^R1763K^* plasmids were grown to mid-logarithmic phase, and identical five-fold serial dilution series were plated on YNB -Trp medium for 3-4 d at 30, 35.5, and 37°C as indicated. (B) Fluorescence microscopy of cells as described in panel *A*, expressing Ste3-GFP. Scale bar, 2 µm. (C) Quantification of endocytic capacity of cells as in panel *B*, but expressing Ste3-pHluorin. Whole-cell fluorescence intensity was measured for n=3 biological replicates, with 47-78 cells measured per condition for each trial. One-way ANOVA with Tukey’s multiple comparisons test, * p<0.05; ** p<0.01.

### Sac7, unlike other RhoGAPs, localizes opposite from sites of Rho1 activity

To exert an effect on endocytosis, GAPs likely need to engage Rho1 and alter its activity at the PM. Localization of RhoGAPs expressed from endogenous promoters has not been comprehensively studied; thus, we generated WT and 4Δ strains expressing GAPs with C-terminal mNeonGreen (mNG) tags (Fig. 4). We found that Bag7-mNG localized to the cytosol and to cytoplasmic punctae, making involvement in CIE unlikely. The other three GAPs localized to cytosol and PM, albeit with different cortical patterns. Bem2-mNG localized preferentially to the bud neck and to a lesser extent at cortical foci on both the mother and daughter side, while Lrg1-mNG localized prominently to the bud tip and bud neck, but not to the mother (Breker et al., 2013). In contrast, Sac7-mNG did not localize to the bud neck, but formed cortical foci that were more prominent on the mother than on the bud. Disruption of CME did not affect RhoGAP localization, as each protein showed similar distribution in WT, 4Δ^ON^ and 4Δ^OFF^ backgrounds. Importantly, Rho1 activity prominently polarizes to the bud tip (Yamochi et al., 1994; Yoshida et al., 2009), opposite the localization for Sac7, suggesting that Sac7 may inactivate Rho1 at the mother PM, thereby negatively regulating CIE.

**Figure 4:**
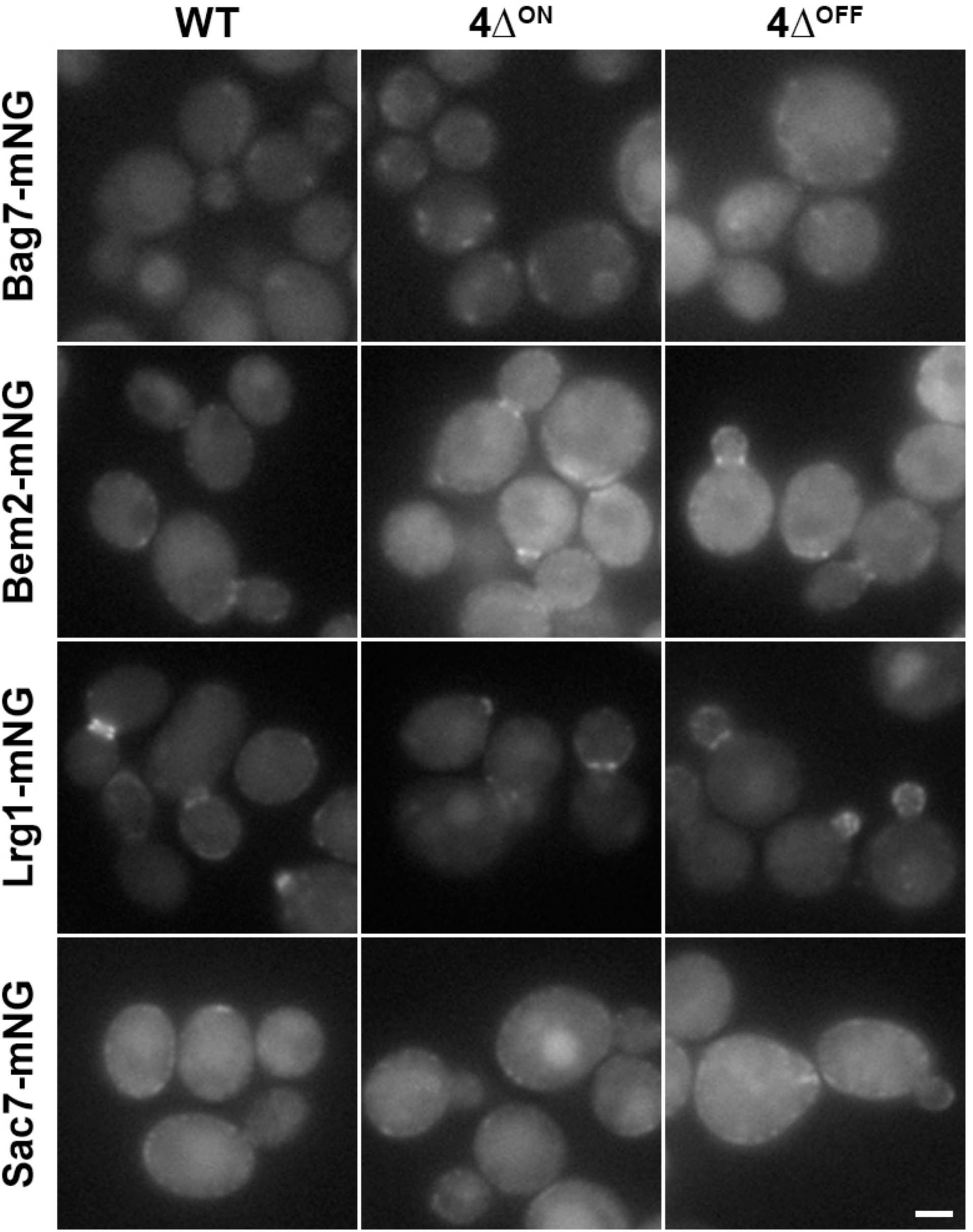
Subcellular localization or Rho1 GAPs. Fluorescence microscopy of WT, 4Δ^ON^ and 4Δ^OFF^ cells expressing Bag7, Bem2, Lrg1, or Sac7 with a C-terminal mNeonGreen tag as indicated. Scale bar, 2 µm.

### Rho1 activity depolarizes in sac7Δ cells

Based on our results, we reasoned that GAP deletion may alter localized Rho1 activity at the PM, and thereby influence CIE. Since GFP-tagged Rho1 cannot distinguish between active, GTP-bound and inactive, GDP-bound states, we generated a fluorescent probe based on Pkc1 as an effector that binds Rho1-GTP. Pkc1 has two Rho-binding domains (RBDs); the first RBD in the N-terminal PKC homology region (HR1), has been used as a probe for active Rho1 (Denis and Cyert, 2005; Kono et al., 2012; Onwubiko et al., 2023), but does not bind Rho1-GTP *in vitro* (Logan et al., 2010). Instead, we fused mCherry to the second Rho-binding domain encoded in the cysteine-rich C1 region (amino acids 384-632, hereafter referred to as mCherry-RBD) since this region biochemically associates with Rho1-GTP (Logan et al., 2010). We assessed localization of our mCherry-RBD fusion in cells co-expressing GFP-Rho1^WT^ or constitutively-active GFP-Rho1^Q68H^ (Supplementary Fig. S1A). As reported previously, GFP-Rho1^WT^ localized to the PM and to internal structures including the vacuole and nuclear membranes (Yoshida et al., 2009). Consistent with function at polarized growth sites, cortical GFP-Rho1^WT^ localization biased toward the tip of budded cells, but was evenly distributed in unbudded cells. GFP-Rho1^Q68H^ showed similar polarized distribution in budded cells and broad PM localization in unbudded cells, but was not observed on internal structures as seen for wild-type Rho1 (Yoshida et al., 2009). mCherry-RBD closely mimicked the PM distribution of GFP-Rho1^Q68H^ in unbudded and budded cells, and similarly did not localize to internal membranes. Instead, a portion of mCherry-RBD was found within the nucleus, which is a known localization site for Pkc1 (Denis and Cyert, 2005). Moreover, we verified that mCherry-RBD does not exert a dominant-negative effect, since its expression does not impact CIE or temperature-dependent growth phenotypes described above (Supplementary Figs. S1B and S2). Overall, our mCherry-RBD acts as an *in vivo* sensor for active Rho1, at least at the PM, similar to other RBD probes (Kono et al., 2012; Onwubiko et al., 2023).

With the mCherry-RBD probe validated, we next assessed its PM localization in WT, 4Δ^ON^ and 4Δ^OFF^ cells with or without RhoGAP deletions (Fig. 5A). As expected, mCherry-RBD preferentially localized to the tip of budding cells consistent with polarized Rho1 activity; this distribution appeared similar in WT, 4Δ^ON^ and 4Δ^OFF^, suggesting that blocking CME does not alter Rho1 polarization. Similar mCherry-RBD distribution was observed in all *bag7*Δ, *bem2*Δ, and *lrg1*Δ strains. Distinct from the other GAPs, *sac7*Δ resulted in prominent mCherry-RBD accumulation on the mother PM. As the only GAP that showed preferential mother PM localization, Sac7 may thus curb mother Rho1 activity, thereby enhancing polarization of active Rho1.

**Figure 5:**
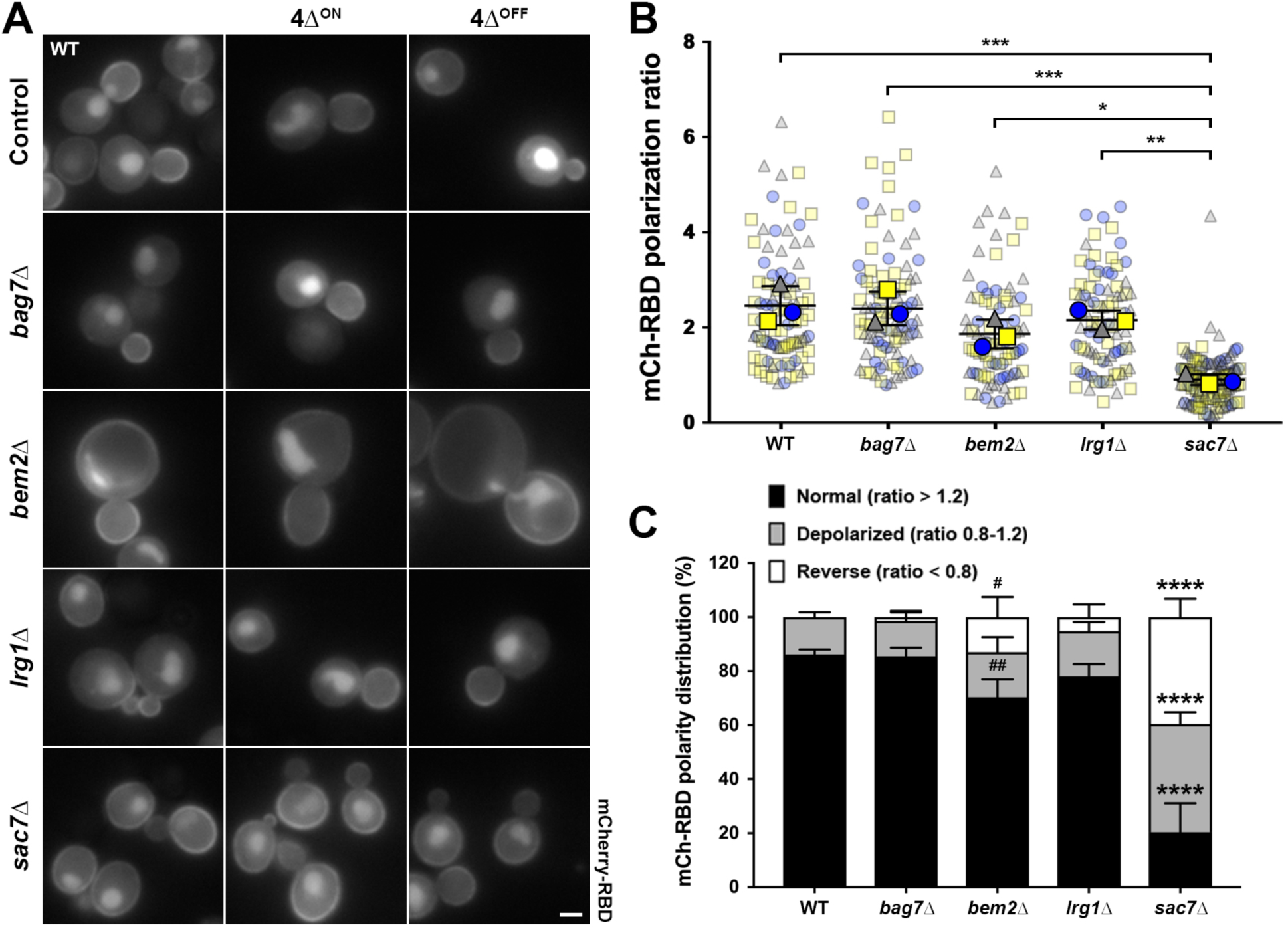
Effect of Rho1 GAP deletion on polarization of Pkc1 Rho-binding domain (RBD), a probe for active Rho1. (A) WT (control), 4Δ^ON^ and 4Δ^OFF^ cells without and with Rho1 GAP deletions as indicated, and expressing mCherry-RBD from a low-copy plasmid, were grown in YNB -Lys -Met medium and imaged by fluorescence microscopy. Scale bar, 2 µm. (B) Quantification of mCherry-RBD polarization in WT, *bag7*Δ, *bem2*Δ, *lrg1*Δ and *sac7*Δ cells. Ratio of peak intensity at the bud tip versus the base of the mother cell was determined from line scans. Polarization ratios were calculated for n=3 biological replicates, with 23-56 cells measured per condition for each trial. One-way ANOVA with Tukey’s multiple comparisons test, * p<0.05; ** p<0.01; *** p<0.001. (C) Classification of WT, *bag7*Δ, *bem2*Δ, *lrg1*Δ, and *sac7*Δ cells with normal polarization (ratio > 1.2), depolarized (ratio between 0.8-1.2), or reverse polarization (ratio < 0.8) of mCherry-RBD. Cells from each trial in panel B were categorized based on the above ratios and expressed as a percentage (n=3). Two-way ANOVA with Tukey’s multiple comparisons test, # p<0.05 and ## p<0.01 compared to the equivalent category in WT and *bag7*Δ; **** p<0.0001 compared to the equivalent category in all conditions. An expanded data figure comparing mCherry-RBD polarization between WT, 4Δ^ON^, and 4Δ^OFF^ cells in each control and Rho1 GAP deletion background is provided in Supplementary Figure S3.

To quantify mCherry-RBD polarization, we measured the ratio of fluorescence intensity at the bud tip to the mother base, and compared WT to RhoGAP deletion strains (Fig. 5B). Unpolarized Rho1 gives a ratio close to 1.0, while polarization toward the bud yields a value greater than 1.0 and reversed polarization toward the mother gives a lower ratio. WT, *bag7*Δ, *bem2*Δ, and *lrg1*Δ cells had RBD polarization ratios near or greater than 2, indicating that Rho1 activity enriched toward the bud tip. In contrast, *sac7*Δ cells had a significantly reduced RBD polarization ratio (0.901 ± 0.114) compared to all other conditions, confirming that loss of Sac7 redistributes Rho1 activity toward the mother. As a separate representation of altered mCherry-RBD polarization, we grouped cells analyzed in Fig. 5B into three RBD polarity distributions: normal (ratio > 1.2, which comprised 86.18 ± 1.83% of WT cells), unpolarized (ratio between 0.8-1.2, which included the remaining 13.82 ± 1.83% of WT cells), and reversed (ratio < 0.8; no WT cells showed this phenotype) (Fig. 5C, distribution examples in Supplementary Fig. S3). Using these classifications, we found that *bag7*Δ and *lrg1*Δ were indistinguishable from WT, while *bem2*Δ gave a small increase in cells with reversed RBD polarity and a concomitant decrease in normal polarization compared to WT. In contrast, *sac7*Δ resulted in a large decrease in normal polarization (to 20.35 ± 10.73%), with a corresponding increase in depolarized (40.10 ± 4.27%) and reversed RBD polarity (39.55 ± 6.74%) compared to WT and all other RhoGAP deletions.

In analyzing mCherry-RBD distribution, we considered whether CME disruption alters active Rho1 polarization. We used the same dataset to compare RBD polarization ratio (Supplementary Fig. S5A-E) and distribution into normal, depolarized, and reverse polarity classes (Supplementary Fig. S5F-J) in WT, 4Δ^ON^ and 4Δ^OFF^ backgrounds. 4Δ^ON^ and 4Δ^OFF^ strains had indistinguishable RBD polarization ratios compared to their WT or RhoGAP deletion counterparts, including in *sac7*Δ cells which showed low RBD polarization regardless of CME status. RBD polarity distributions revealed that 4Δ^ON^ and 4Δ^OFF^ only had minor differences for *bag7*Δ and *lrg1*Δ that are likely unrelated to CIE (Fig. S4G-I), since neither gene affected cargo internalization in 4Δ^OFF^.

A final factor we considered is whether altered RBD distribution in *sac7*Δ results from loss of active Rho1 at the bud tip, or from failure of *sac7*Δ cells to inactivate Rho1 at the mother cortex. We assessed mCherry-RBD peak intensity at the bud tip or mother base (Supplementary Figs. S5A-B) for cells examined in Fig. 5B. If *sac7*Δ depolarizes RBD by reducing Rho1 activity at the bud tip we would expect lower bud tip intensity, but did not observe any difference compared to WT. The alternative of *sac7*Δ reducing Rho1 inactivation and extraction from the mother cortex was supported by our finding that RBD intensity increase at the mother base in *sac7*Δ. *BEM2* deletion increased RBD intensity at the bud tip and mother base compared to other conditions, even though polarization was similar to WT, which might result from *bem2*Δ cell and vacuolar size defects or compression of cytosolic mCherry-RBD toward the cell periphery (Fig. 1B) (Marquitz et al., 2002). Taken together, our results indicate that *sac7*Δ reduces Rho1 inactivation and removal from the mother cortex without altering bud tip activation.

### sac7Δ increases Bni1 retention at non-CME cortical patches

Given the increased cortical RBD levels in *sac7*Δ, we reasoned that elevated PM Rho1-GTP might enhance effector recruitment for downstream signaling. Since Bni1 is required for Rho1-dependent CIE (Prosser et al., 2011), we examined Bni1-3GFP localization and dynamics in WT and RhoGAP-deficient cells. Because 4Δ^ON^ and 4Δ^OFF^ did not considerably impact mCherry-RBD distribution (Supplementary Fig. S4), we limited our analysis to cells expressing all four endocytic adaptors. Previous studies showed that Bni1 localizes to the bud tip and neck, and to highly dynamic cytoplasmic speckles (Buttery et al., 2007). To visualize these structures, we performed time-lapse imaging using near-TIRF (total internal reflection fluorescence) microscopy (Tokunaga et al., 2008). In budding cells, we observed the expected bud tip and/or neck localization of Bni1-3GFP, as well as punctae that moved rapidly within the cytoplasm in WT and RhoGAP deletion strains (Fig. 6A and Movies 1-5). Additionally, we occasionally observed static cortical Bni1-3GFP punctae that persisted for several seconds and sometimes for the entire movie (Fig. 6A, yellow arrowheads). These stabilized cortical Bni1 foci were more prevalent in *sac7*Δ cells and predominantly localized to the mother.

**Figure 6:**
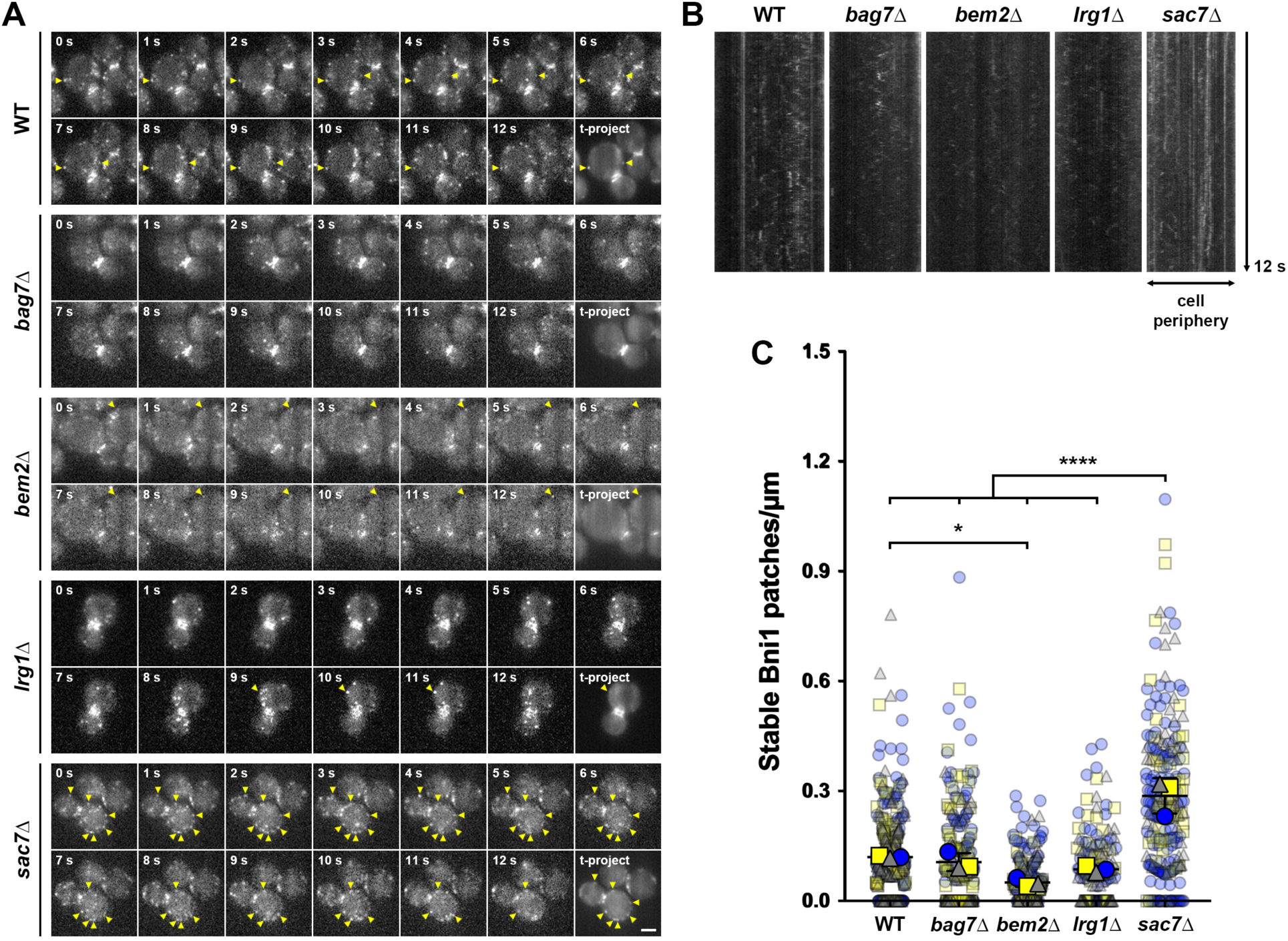
Effect of *SAC7* deletion on cortical recruitment and dynamics of Bni1. (A) Near-TIRF, timelapse imaging of WT, *bag7*Δ, *bem2*Δ, *lrg1*Δ, and *sac7*Δ cells expressing Bni1-3GFP was performed at 20 fps for 12 s. Bleach-corrected still frames are shown at one-second intervals as indicated, while the last image panel for each strain consists of an average intensity time projection (t-project) of the entire movie to identify stabilized patches (yellow arrowheads, also indicated on each 1 s interval frame that contains a patch visible in the time projection). Images shown are stills from Movies 1-5. Scale bar, 2 µm. (B) Kymographs from cells imaged as in panel A were generated using the KymoResliceWide plugin in Fiji. (C) Quantification of stabilized Bni1-3GFP cortical patches in WT, *bag7*Δ, *bem2*Δ, *lrg1*Δ, and *sac7*Δ cells. Bni1-3GFP patches per µm of cell perimeter were counted in time projection in n=3 biological replicates, with 32-156 cells measured per condition for each trial. One-way ANOVA with Tukey’s multiple comparisons test, * p<0.05; **** p<0.0001.

We took several approaches to verify Bni1 patch stabilization in *sac7*Δ cells. First, we generated kymographs from lines drawn around the mother periphery, which showed increased static traces indicative of PM stabilization in *sac7*Δ compared to WT and other GAP deletion strains (Fig. 6B). Second, we generated average intensity projections from timelapse series, effectively treating the data as a three-dimensional stack (x:y:time instead of x:y:z; Fig. 6A). In these “time-projections”, mobile speckles fade into the background while stabilized structures appear as distinct patches. We observed infrequent stabilized Bni1 patches at the cortex of WT, *bag7*Δ, *bem2*Δ, and *lrg1*Δ cells, with an increase in *sac7*Δ. Lastly, we quantified density of stabilized cortical Bni1-3GFP patches from time-projections (Fig. 6C). When corrected for cell perimeter, we found that *bem2*Δ had lower stable patch density compared to WT, likely due to increased cell size. In contrast, *sac7*Δ resulted in significantly higher Bni1-3GFP patch density compared to WT and other GAP-deficient strains.

In yeast, cortical actin patches are CME sites in which Arp2/3 stimulates branched actin network assembly to support membrane bending (Kaksonen et al., 2003). Since formins elongate unbranched actin filaments that are mutually exclusive from the Arp2/3 network (Burke et al., 2014), and because most proteins implicated in CIE do not localize to CME sites, we predicted that stabilized Bni1 patches would be distinct from cortical actin patches. Accordingly, we generated time-projections as described in Fig. 6, but with cells co-expressing Bni1-mNeonGreen and Abp1-mCherry as a marker for CME sites. As with Bni1-3GFP, Bni1-mNeonGreen showed increased stable patches in *sac7*Δ compared to WT or other RhoGAP deletions (Fig. 7). Notably, stable Bni1-mNeonGreen patches did not co-localize with Abp1-mCherry cortical patches, indicating that these sites are distinct from CME.

**Figure 7:**
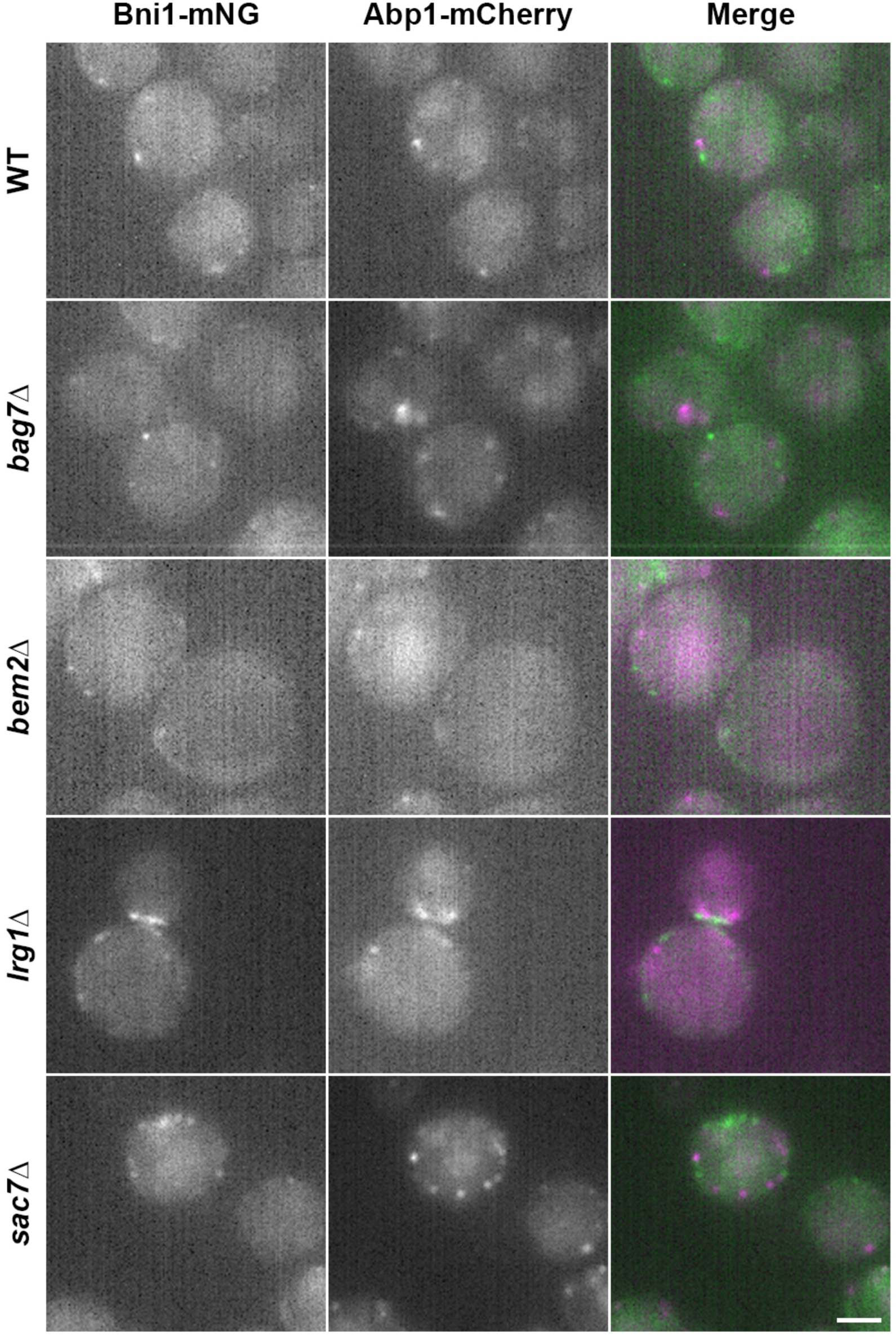
Distinction of stable Bni1 sites from cortical actin patches. Simultaneous two-color, near-TIRF timelapse imaging of WT, *bag7*Δ, *bem2*Δ, *lrg1*Δ, and *sac7*Δ cells expressing Bni1-mNeonGreen (green) and Abp1-mCherry (magenta) was performed at 20 fps for 12 s. Image series were then beach corrected prior to generating average intensity time projections. Scale bar, 2 µm.

### sac7Δ alters actin morphology

Since Bni1 promotes actin elongation, we hypothesized that increased cortical retention in *sac7*Δ would alter actin morphology. Whereas Abp1 associates with Arp2/3-generated branched actin networks at cortical patches, Abp140 associates with actin cables and unbranched filaments (Huckaba et al., 2004). To examine effects of RhoGAP deletion on actin morphology, we generated strains expressing Abp140-3GFP and Abp1-mCherry from endogenous promoters and created projection images from deconvolved z-stacks (Fig. 8A). In WT cells, Abp140-3GFP formed actin cables that oriented along the mother/bud axis; these cables were more prominent in the mother than in the bud, as reported previously (Wirshing and Goode, 2024). Abp1-mCherry formed patches distinct from, but often proximal to, Abp140-3GFP cables, with greater density in the bud consistent with higher endocytic rates in daughter cells (Huckaba et al., 2004). Similar cable and patch morphologies and distributions were observed in RhoGAP deletions, in particular *bag7*Δ and *lrg1*Δ. Cables were more readily observed in the buds of *bem2*Δ cells, and appeared slightly less oriented along the polar axis, although it is unclear whether increased cell size accounted for these differences. In *sac7*Δ, Abp140-3GFP cables oriented toward the bud, but we additionally observed Abp140 varicosities or patch-like structures that were not obviously associated with longer actin cables and did not co-localize with Abp1-mCherry. It is unclear whether these represent shorter cables or altogether different structures; however, quantification of Abp140-3GFP patch-like formations revealed they were present at nearly 3-fold higher density in *sac7*Δ compared to any other condition (Fig. 8B). These data suggest that Sac7 influences actin cable morphology, while the other RhoGAPs may play less prominent roles.

**Figure 8:**
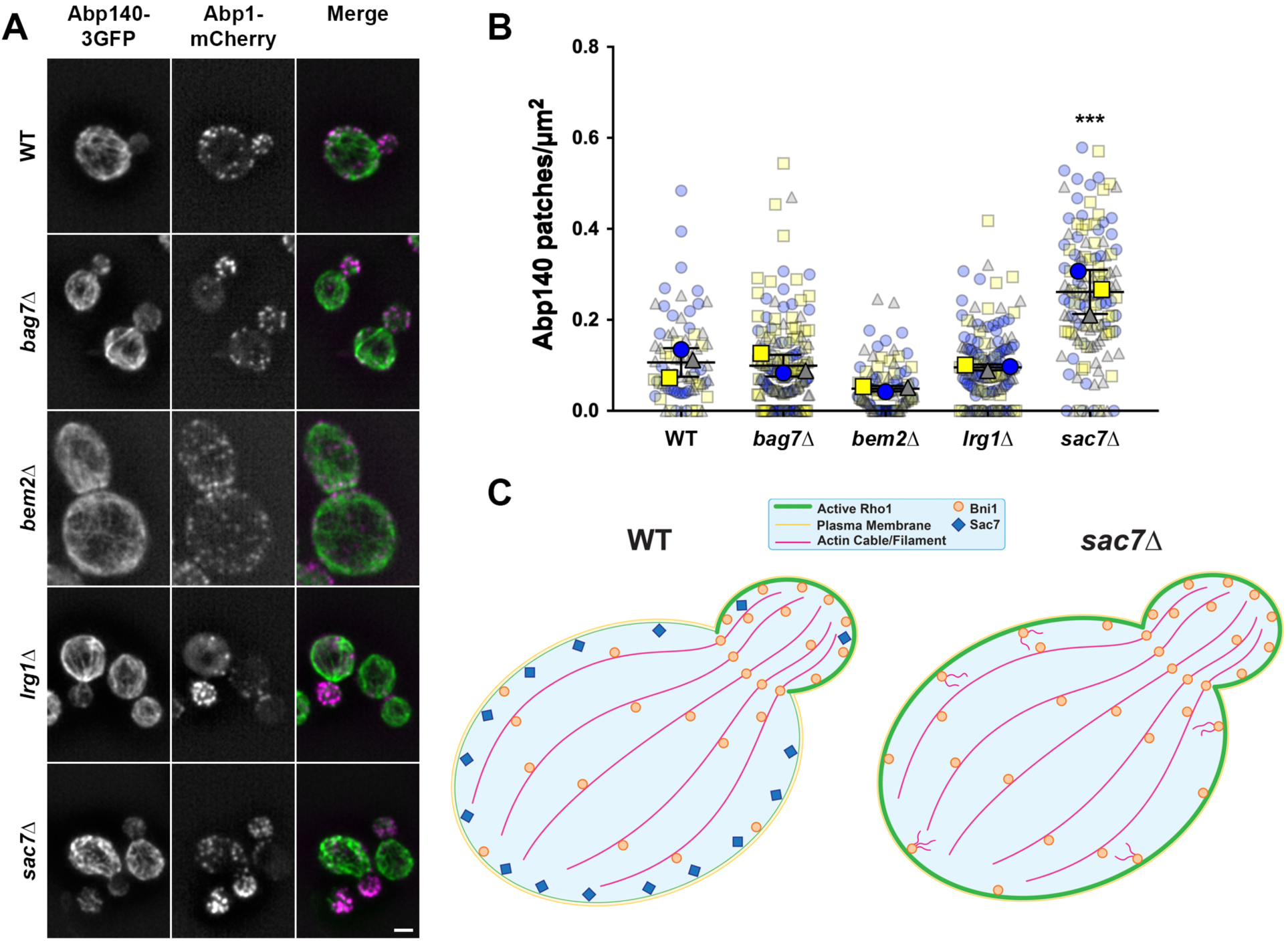
Effect of Rho1 GAP deletion on actin morphology, and model for the role of Sac7 in regulating Rho1 localization and clathrin-independent endocytosis. (A) Z-stacks of WT, *bag7*Δ, *bem2*Δ, *lrg1*Δ, and *sac7*Δ cells expressing Abp140-3GFP (green) and Abp1-mCherry (magenta) were collected at 0.2 µm step intervals. Stacks were then deconvolved and converted into summed z-projections. Scale bar, 2 µm. (B) Quantification of patch-like Abp140-3GFP structures, corrected for cell size, in WT and RhoGAP deletion strains. n=3 biological replicates, with 15-72 cells measured per condition for each trial. One-way ANOVA with Tukey’s multiple comparisons test, *** p<0.001 compared to all other conditions. (C) Sac7 localizes preferentially to the mother cortex in WT cells, where it locally inactivates Rho1 (left). In contrast, active Rho1-GTP localizes to the bud and facilitates Bni1 recruitment for generation of polarized actin cables. In *sac7*Δ cells (right) Rho1-GTP accumulates at the mother cell cortex, leading to depolarized Rho1 activity. Elevated cortical Rho1-GTP increases Bni1 recruitment at the plasma membrane, resulting in altered actin polymerization which may facilitate CIE.

## Discussion

Mechanisms governing Rho1 activity related to endocytosis are undercharacterized. Here, we demonstrate that the RhoGAP Sac7 contributes to CIE, and our results showing that *sac7*Δ restores cargo internalization in CME-deficient cells suggest that Sac7 is a key player in yeast CIE. To our knowledge, this is the first example of a GAP playing a negative regulatory role in clathrin-independent internalization. Studies of mammalian CIE showed that expression of the RhoA and Cdc42 GAP Graf1 promotes clathrin-independent carrier (CLIC) formation at the PM, suggesting RhoGAP regulation of endocytosis is not restricted to yeast (Francis et al., 2015; Lundmark et al., 2008). Importantly, numerous CIE pathways rely on Rho, Arf, and Ras family small GTPases, suggesting that positive and/or negative roles for GAPs in shaping localized GTPase activity during endocytosis may be more prevalent than we currently realize.

Our current study builds on the detailed CIE framework from our past studies in yeast as a genetically tractable system to define endocytic factors, in which we identified the GEF Rom1, its PM recruitment factor Mid2, and the Rho1 GTPase as key facets of CIE (Prosser et al., 2011). Downstream, the formin Bni1 and stabilizers of unbranched actin filaments are necessary for clathrin-independent cargo internalization. Further, we identified type V myosin (Myo2), cytoplasmic microtubules and dynein/dynactin, and cargo-selective α-arrestins as contributors to CIE (Prosser et al., 2015; Woodard et al., 2023). Together, these findings lead us to our current mechanistic model for CIE, where GEFs and GAPs collectively regulate Rho1 PM activity and signaling through its effector Bni1 to control actin assembly needed for membrane internalization (Fig. 8C).

With two endocytic routes defined in yeast, how are cargos sorted between them? This remains an open question in the field, but there are some hints of proteins such as α-arrestins and Syp1 contributing to both pathways, albeit through different mechanisms (Apel et al., 2017; Prosser et al., 2015). Notably, a previous study described roles for Rho1 and Sac7 in mating pheromone uptake which also required the CWI sensor Wsc1 and the Rho1 effector Fks1, neither of which are involved in CIE (deHart et al., 2003; Prosser et al., 2011). In contrast, Mid2 and Bni1, both involved in CIE, were not required for pheromone uptake (deHart et al., 2003). The study was published prior to our initial characterization of yeast CIE, and did not assess whether Rho1 and Sac7 acted with or separate from CME machinery. The current study demonstrating CIE activity for Rho1 and Sac7 raises the possibility of cross-talk between CME and CIE pathways, which will be further explored in future studies.

Whereas GEFs are known to promote localized GTPase recruitment and activation, the relative contribution of GAPs is less understood but may be important for restricting activity and engagement with downstream effectors at sites where function is unnecessary. Spatiotemporal activity of small GTPases is tightly regulated to ensure precise vesicular targeting and organelle identity (Rab and Arf), signal transduction (Ras), and actin architecture (Rho), and to prevent inappropriate activation detrimental to cell function. Such functional regulation is achieved through the complex interplay between activation and inactivation cycles respectively mediated by GEFs and GAPs, membrane insertion and extraction via GDP-dissociation inhibitors (GDIs), and upstream events initiated by receptor signaling and recruitment factors. In yeast, Rho1 and Cdc42 localize to the PM and on internal organelles, but the GTP-bound active state strongly polarizes toward sites of growth (bud tip or mating projection) (Lu and Drubin, 2022; Watson et al., 2014; Yoshida et al., 2009). Both proteins regulate cell polarity and formin-dependent assembly of actin cables that enable secretory vesicle and organelle delivery to growth sites (Evangelista et al., 1997; Evangelista et al., 2001; Pruyne et al., 2002; Yu et al., 2011). Delivery and GEF-mediated activation of Rho1/Cdc42 at growth sites is critical for polarity establishment, while secretory vesicle fusion and lateral diffusion of prenylated GTPases anchored to the PM inner leaflet create an activity gradient (Watson et al., 2014; Woods et al., 2015; Woods et al., 2016). Actin, endocytosis, and endocytic recycling may also contribute to Cdc42 polarization (Freisinger et al., 2013; Marco et al., 2007; Watson et al., 2014); however, Rho1 internalization from the PM has not been documented in yeast to our knowledge and it is unclear whether Rho1 is an endocytic cargo in addition to its role in promoting CIE. Since the bud neck represents a diffusion barrier, GAP localization to this site may contribute to tightening activity gradients (Luedeke et al., 2005). Indeed, the Rho1 GAPs Bem2 and Lrg1 (Fig. 4), and the Cdc42 GAPs Rga1 and Bem3 (Breker et al., 2013), localize to the bud neck where they may additionally prevent premature Rho1/Cdc42 activity and contractile actin assembly at the cytokinesis site.

Our study revealed that Sac7 PM distribution at the mother cortex inversely correlates with Rho1-GTP levels assessed using mCherry-RBD as a probe for active Rho1 (Fig. 4 and 5A). This opposite localization suggests Sac7 restricts Rho1 function at the mother PM, tightening the polar gradient (Fig. 8C). The observed depolarization of active Rho1 in *sac7*Δ has functional implications on its effector Bni1, which shows enhanced recruitment or retention at the cell cortex. Subsequently, Bni1 may alter actin cable morphology or unbranched filament polymerization to provide forces necessary for membrane bending during CIE. Whether negative regulation of CIE is a primary function of Sac7 is unclear, as its role in regulating both actin polymerization and CWI signaling suggests a role in re-establishing actin cable polarization after cell wall repair (Schmidt et al., 2002). To our knowledge, a role for endocytosis in actin repolarization after cell wall damage has not been assessed. Moreover, *sac7*Δ results in a cold-lethal phenotype that has not been extensively studied, but may arise due to inefficient Rho1-GTP polarization or actin cable assembly (Dunn and Shortle, 1990). Even though *sac7*Δ eliminates the active Rho1 gradient, it likely is not lethal because essential Rho1 activity remains, and because localization and activation of Cdc42, the master regulator of yeast cell polarity, appears independent of Rho1. Whether and how *sac7*Δ impacts temperature-dependent CWI signaling, Rho1 activity, actin organization, and CIE is unknown, and will be the subject of future studies.

Non-overlapping subcellular localization of yeast RhoGAPs is consistent with distinct cellular roles or downstream effector regulation. Our results suggest that GAPs shape active Rho1 localization to impact downstream functions, including actin morphology. Since Rho1 has multiple effectors, it will be interesting to determine whether GAP loss-of-function also impacts localization or function of other downstream binding partners. Notably, RhoGAP-mediated negative regulation is observed outside of yeast. For example, RhoA, Rac1 and Cdc42 regulate actin-dependent growth cone guidance in neurons, and localized function of the GAPs p190 and ARHGAP4 restricts growth cone extension (Brouns et al., 2001; Foletta et al., 2002; Koinuma et al., 2017; Vogt et al., 2007). Similarly, Myo9b GAP activity toward RhoA, and its Rac1-dependent recruitment to the leading edge of leukocytes, locally restricts RhoA-GTP levels that impair migration (Hemkemeyer et al., 2020). Our future work will identify localization determinants and binding partners for RhoGAPs to better understand how they regulate endocytic and signaling events.

## Materials and Methods

### Reagents, yeast strains, and plasmids

All yeast strains used in this study were generated in the SEY6210 genetic background, and are described in Supplementary Table 1, while plasmids are described in Supplementary Table 2. Unless otherwise noted, cells were grown at 30°C in YPD (yeast extract, peptone, dextrose) medium or in YNB (yeast nitrogen base; synthetic defined) medium lacking uracil, tryptophan and/or lysine as needed for plasmid selection, and lacking methionine in experiments inducing expression of genes controlled by the *MET25* promoter. Gene deletions and C-terminal, cassette-based fluorescent protein tagging were conducted using PCR-based genomic integration strategies as described previously (Goldstein and McCusker, 1999; Longtine et al., 1998). All yeast transformations were performed using the lithium acetate method (Schiestl and Gietz, 1989).

Strains expressing *ABP140-3GFP::LEU2* were generated by transformation of plasmid pB1994 linearized with NdeI, followed by selection on YNB medium lacking leucine. Strains expressing *BNI1-3GFP::LEU2* were similarly generated by transformation of plasmid pB1993 linearized with BssHI, followed by selection on YNB medium lacking leucine. pB1993 and pB1994 were kindly provided by Dr. David Pellman (Harvard University) (Buttery et al., 2007).

Plasmid pDP0551 was generated by PCR amplification of the Pkc1 Rho-binding domain (amino acids 384-632) from SEY6210 genomic DNA and cloning into pUG34-mCherry to fuse the Rho-binding domain to an N-terminal mCherry tag under control of the *MET25* promoter, which drives expression in the absence of methionine (Mumberg et al., 1994; Sangsoda et al., 1985), and a *CYC1* terminator. The tagged construct with promoter and terminator was then amplified using T3 and T7 primers, and subcloned by homologous recombination into pRS317 linearized with EcoRI and NotI.

Plasmids pDP0480 and pDP0481, expressing GFP-Rho1^WT^ and GFP-Rho1^Q68H^, respectively, were generated by PCR amplification of the coding sequence plus endogenous promoter and terminator from plasmids SP301 and SP307 (kindly provided by Dr. David Pellman) using M13 forward and reverse primers (Yoshida et al., 2009). Amplified sequences were then subcloned into pRS317 linearized with EcoRI and NotI.

Unless otherwise indicated, reagents were purchased from Sigma-Aldrich (St. Louis, MO) or Thermo Fisher Scientific (Waltham, MA). Restriction enzymes were purchased from New England Biolabs (Ipswitch, MA).

### Plate-based growth assays

To assess differences in temperature-dependent growth, cells were inoculated into liquid YPD medium, or in YNB medium lacking tryptophan for selection of *SAC7* expression plasmids, and grown overnight in a 30°C shaking incubator. Selection for Ent1- or ENTH1-expressing plasmids in the 4Δ background was not necessary because loss of the plasmid in these strains is lethal (Aguilar et al., 2006; Maldonado-Báez et al., 2008; Prosser et al., 2011). Cells were then diluted in fresh medium to an OD_600_ of 0.3-0.35, and re-grown at 30°C until they reached mid-logarithmic phase (∼0.6-0.8 OD_600_). Culture densities were adjusted to an OD_600_ of 0.25, and cells were subjected to a five-fold serial dilution series (1:1, 1:5, 1:25). Equal volumes of each dilution were transferred to a sterile 32-well circular dish, and cells were spotted onto YPD or YNB minus tryptophan plates using a replicative pronging device (Dan-Kar Corp., Woburn, MA). Triplicate plates were grown at 30, 35.5, and 37°C for 3-4 d before imaging using a FluorChem M gel documentation system (Bio-Techne, San Jose, CA).

### Fluorescence microscopy and post-acquisition image analysis

All microscopy was performed using a DMi8 inverted fluorescence microscope (Leica Microsystems, Deerfield, IL) equipped with an infinity TIRF module, a 100X, 1.47 NA Plan-Apochromat oil immersion objective, a 1-1.6X variable optovar, an LED3 fluorescence illumination system, excitation/emission filter cubes for GFP and/or mCherry detection, a Flash 4.0 v3 sCMOS camera (Hamamatsu, Bridgewater, NJ), and LAS X v3.7.6.25997 software (Leica). For simultaneous two-color imaging, the system was additionally equipped with a W-View Gemini image splitting device (Hamamatsu), 488 and 561 nm lasers, and a Quad widefield (QWF-S-T) filter cube.

Imaging for all experiments was performed at room temperature using live cells that were grown to mid-logarithmic phase at 30°C. Within an experiment, random fields of cells were chosen for imaging using brightfield mode, without looking at the fluorescence signal, to avoid selection bias and to reduce photobleaching prior to image acquisition. All images within an experiment were collected using identical acquisition parameters to permit direct comparison between strains.

Post-acquisition image analysis was performed using either LAS X for 3D deconvolution, or Fiji/ImageJ2 v2.9.0/1.53t for all other analyses. Experiment-specific procedures are described in the sections below. Within any experiment, all images analyzed were subjected to identical data analysis workflows. Inclusion/exclusion criteria for all cells were pre-determined as described previously (Prosser et al., 2016).

### Quantification of endocytic capacity using Ste3-pHluorin

To quantify changes in endocytic capacity, cells expressing genetically-encoded *STE3-pHluorin* were imaged by fluorescence microscopy, and whole-cell fluorescence intensity was measured as described previously (Prosser et al., 2010; Prosser et al., 2016). Briefly, 16-bit images were background subtracted in Fiji using a 50-pixel rolling ball radius, and regions of interest (ROIs) for individual cells were outlined using the freehand selection tool. Mean gray value (integrated density/area) was then measured for all selected cells to assess fluorescence intensity, corrected for cell size. Experiments were performed as biological triplicates, where all images for a replicate experiment were collected on the same day.

### Quantification of mCherry-Pkc1^RBD^ polarization

The Rho-binding domain of Pkc1 (amino acids 384-632) bearing an N-terminal mCherry tag and expressed from a *MET25* promoter was cloned into pRS317 [*CEN LYS2*] as described above. Transformed WT and RhoGAP deletion cells were selected on YNB medium lacking lysine, and expression of the mCherry-RBD was induced by overnight growth of cells on YNB medium lacking lysine and methionine. mCherry-RBD was imaged by fluorescence microscopy, and images were background subtracted as described above for Ste3-pHluorin quantification. We restricted our analysis to budding cells because we could not assign a “front” or “back” to unbudded cells. RBD polarization was assessed by drawing a straight line passing through the cell from the bud tip to the base of mother, creating an ROI. Intensity along the line was measured using the Plot Profile tool, and values were exported to Microsoft Excel. Peak intensity values were determined at the point of entry (bud tip) and exit (mother base), and the RBD polarization ratio was calculated as the quotient of [bud tip intensity/mother base intensity]. Experiments were performed as biological triplicates, where all images for a replicate experiment were collected on the same day.

To calculate the distribution of RBD polarity ratios, all cells analyzed within each biological replicate experiment were categorized as having normal polarity (ratio > 1.2), depolarized RBD (polarity ration between 0.8-1.2) or reversed polarity (ratio < 0.8), and the percentage of cells in each category was then calculated.

### Assessment of Bni1-3GFP cortical patch stabilization

Cortical recruitment and stabilization of Bni1-3GFP was assessed by time-lapse fluorescence microscopy in near-TIRF mode (Tokunaga et al., 2008). After calibration of the microscope system to calculate excitation field depths using the infinity TIRF module, laser angles were adjusted just outside of the TIRF range. Movies of WT and RhoGAP deletion cells expressing genetically-encoded *BNI1-3GFP* with or without *ABP1-mCherry* were collected at a rate of 20 fps for 241 frames (12 s total elapsed time after the initial time zero frame). Movies were bleach corrected in Fiji using the histogram matching method, and kymographs were generated from movies by drawing a freehand line around the mother cell perimeter and using the KymoResliceWide plugin. Cells co-expressing Bni1-3GFP and Abp1-mCherry were imaged using a W-View Gemini image splitting device for simultaneous acquisition of green and red fluorescence.

To visualize stabilized Bni1-3GFP patches, bleach-corrected movies were converted to average-intensity projection images (x:y:time instead of x:y:z), where structures that remained static over time appeared as patches, and highly motile cytoplasmic or cortical Bni1 speckles disappeared into the averaged background fluorescence. Stabilized patches were quantified by outlining individual cells with the freehand selection tool to create an ROI, counting the number of patches in time projection images, and calculating patch density corrected for cell perimeter.

### Actin morphology analysis

Morphology of actin cables (labeled with Abp140-3GFP) was analyzed in WT and RhoGAP deletion strains expressing genetically-encoded *ABP140-3GFP* and *ABP1-mCherry*. We collected z-stacks at 0.1 µm step intervals through the entire cell thickness, where red and green fluorescence were collected simultaneously using a W-View Gemini image splitting device. Z-stacks were then deconvolved using 10 iterations of the blind deconvolution algorithm in LAS X without removing background or rescaling intensity, and converted into 32-bit sum projection images in Fiji. The density of Abp140-3GFP patch-like structures that were not clearly associated with actin cables was determined by manual counting and correction for cell size.

### Data transparency and statistical analysis

All quantitative experiments were performed as biological triplicates, with the range of cells analyzed per replicate experiment described in accompanying figure legends. Quantitative graphs were presented as SuperPlots prepared in Prism 7 (GraphPad), where background scatter plots show all cells measured for each biological replicate color- and shape-coded by trial (blue circles, yellow squares, and grey triangles) (Lord et al., 2020). Foreground points with the corresponding color/shape scheme represent the trial-level mean of each biological replicate, while bars represent the overall mean and standard deviation calculated from the three biological replicates. RBD polarity distributions were displayed as stacked bar graphs showing means and standard deviations for each category.

Statistical analyses were performed in Prism 7 using one- or two-way ANOVA, as indicated in figure legends, with Tukey’s multiple comparisons test. Analyses were based on trial-level means, where P-values less than 0.05 were considered statistically significant.

## Acknowledgements

We thank Drs. Allyson O’Donnell (University of Pittsburgh), Anita Manogaran (Marquette University), Jason Newton (Virginia Commonwealth University), Beverly Wendland (West Virginia University), and members of the Prosser, Newton, and Wendland labs for helpful discussions, feedback, and critical comments. Matthew Peck, Mohammad Rizavi, and Kamyar Sharifi provided technical assistance with early phases of the study. We also thank Dr. David Pellman (Harvard University) for sharing GFP-Rho1, Abp140-3GFP, and Bni1-3GFP plasmids.

---

**Supplementary Figure S1:**
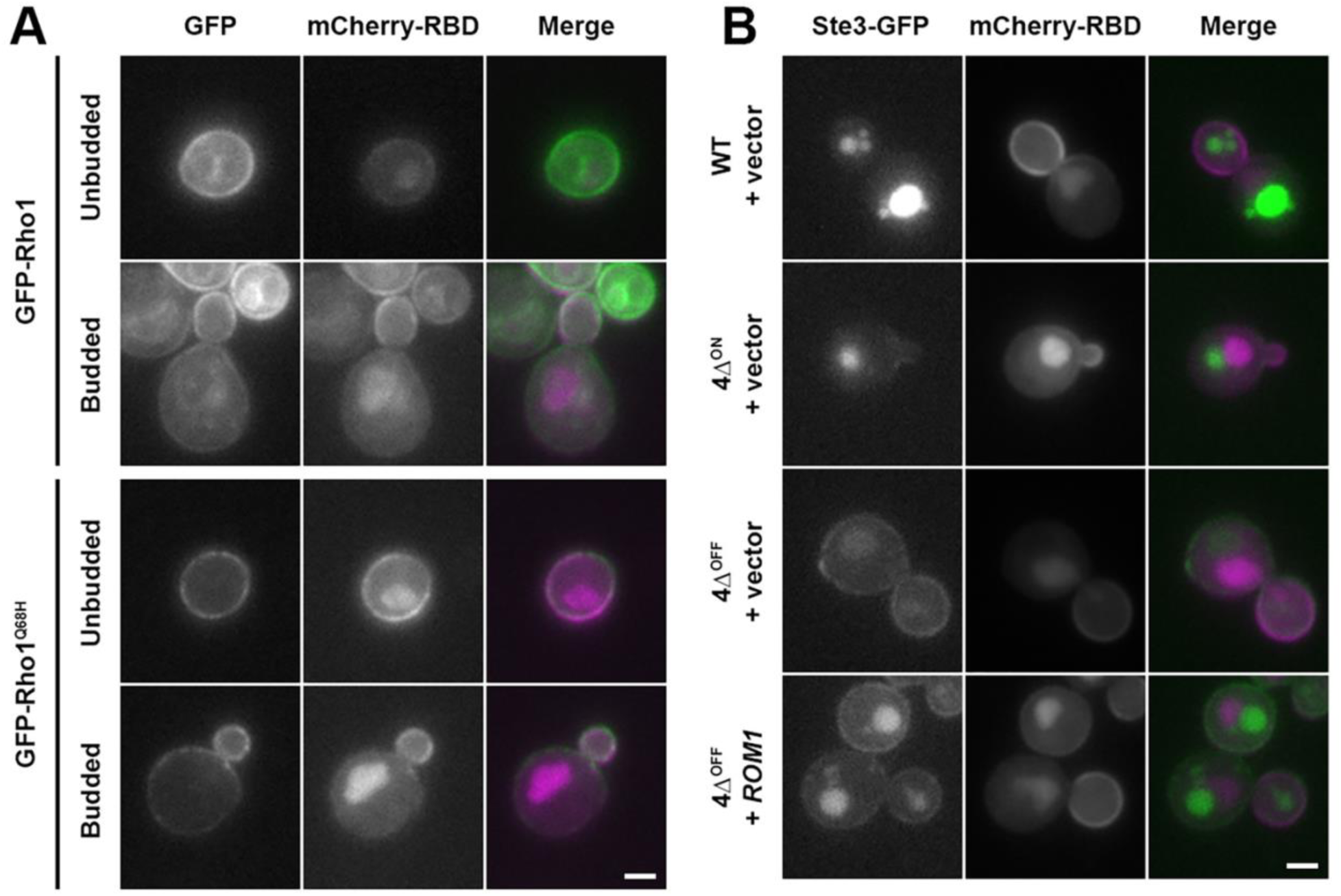
Pkc1^RBD^ as a probe for activated Rho1. (A) Fluorescence microscopy of WT cells transformed with mCherry-RBD and either GFP-Rho1 (upper panels) or GFP-Rho1^Q68H^ (lower panels). Cells were grown on YNB -ura -Lys -Met for plasmid selection and induction of mCherry-RBD expression, and images show unbudded and budded cells as indicated. (B) WT, 4Δ^ON^ and 4Δ^OFF^ cells expressing Ste3-GFP were transformed with empty vector or high-copy *ROM1* [2µ *URA3*] and mCherry-RBD [*CEN LYS2*] plasmids, grown on YNB -ura -Lys -Met for plasmid selection and induction of mCherry-RBD expression, and imaged by fluorescence microscopy. High-copy *ROM1* can still promote Ste3-GFP internalization in 4Δ^OFF^ cells compared to empty vector when mCherry-RBD expression is induced. Scale bars, 2 µm.

**Supplementary Figure S2:**
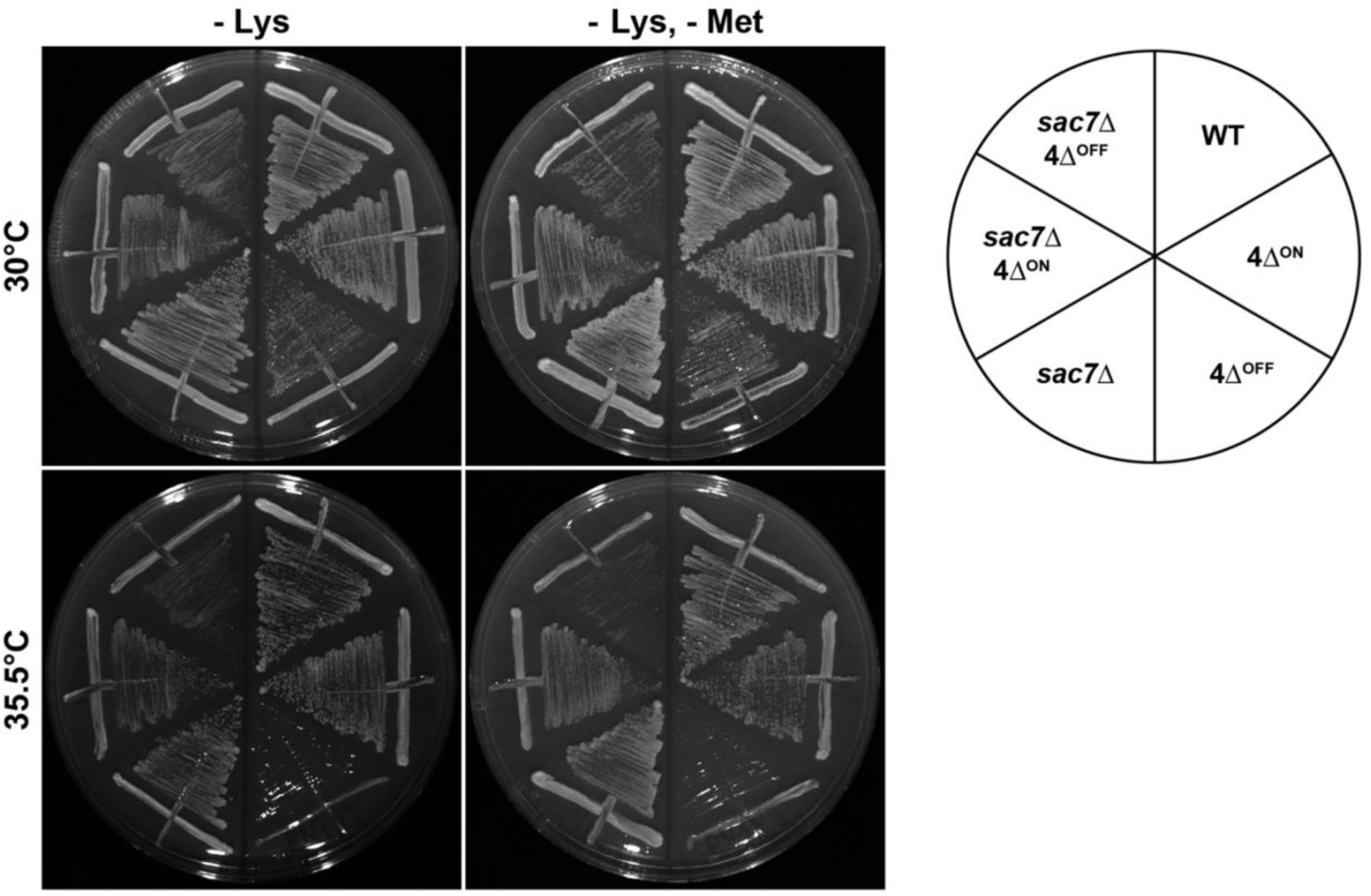
Methionine-induced expression of Pkc1^RBD^ does not alter rescue of temperature sensitivity in *sac7*Δ 4Δ^OFF^ cells. WT, 4Δ^ON^, and 4Δ^OFF^ cells with and without *sac7*Δ were transformed with mCherry-RBD [*CEN LYS2*] under control of the *MET25* promoter, and grown for 3-4 days on YNB -Lys or YNB -Lys -Met at 30 or 35.5°C as indicated. At elevated temperature, *sac7*Δ 4Δ^OFF^ cells grew more robustly than *SAC7* 4Δ^OFF^ cells whether or not mCherry-RBD expression was induced by withdrawal of methionine from the medium.

**Supplementary Figure S3:**
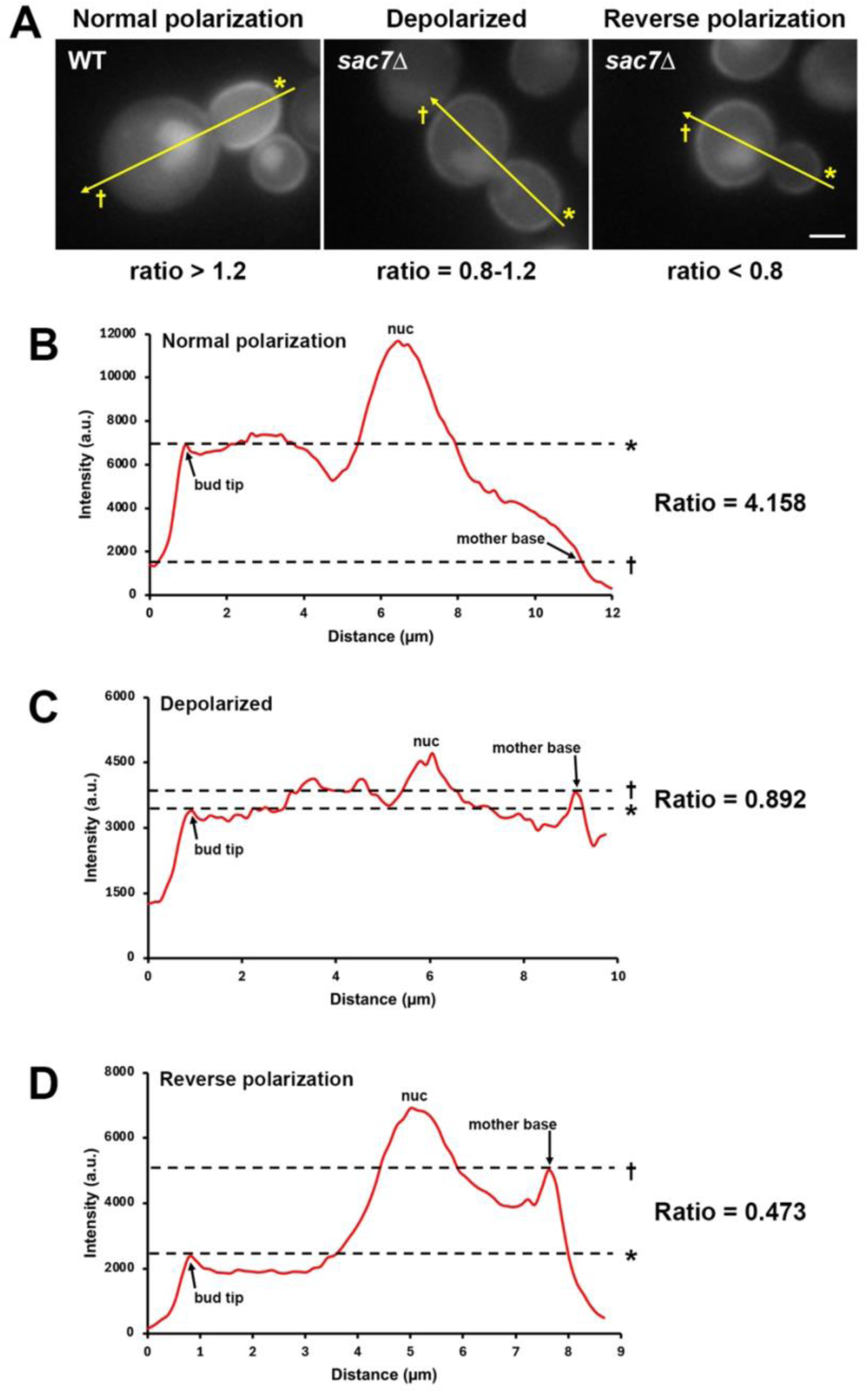
Examples of mCherry-Pkc1^RBD^ polarization and calculation of the RBD polarization ratio. (A) Representative images of WT and *sac7*Δ cells showing normal, depolarized, or reverse polarization of mCherry-RBD as indicated. Yellow arrows correspond to the position of line scans, with the bud tip position marked with an asterisk (*) and the mother base position marked with a dagger (†). Scale bar, 2 µm. (B-D) Line scans from cells shown in panel A, depicting intensity profiles for normal (B), depolarized (C), and reverse polarization (D) of mCherry-RBD. Horizontal dashed lines correspond to the peak intensity at the bud tip (*) or mother base (†), with RBD polarization calculated as the ratio of bud tip to mother base intensity. Note that intensity peaks at the center of the line scan correspond to nuclear localization of mCherry-RBD (nuc).

**Supplementary Figure S4:**
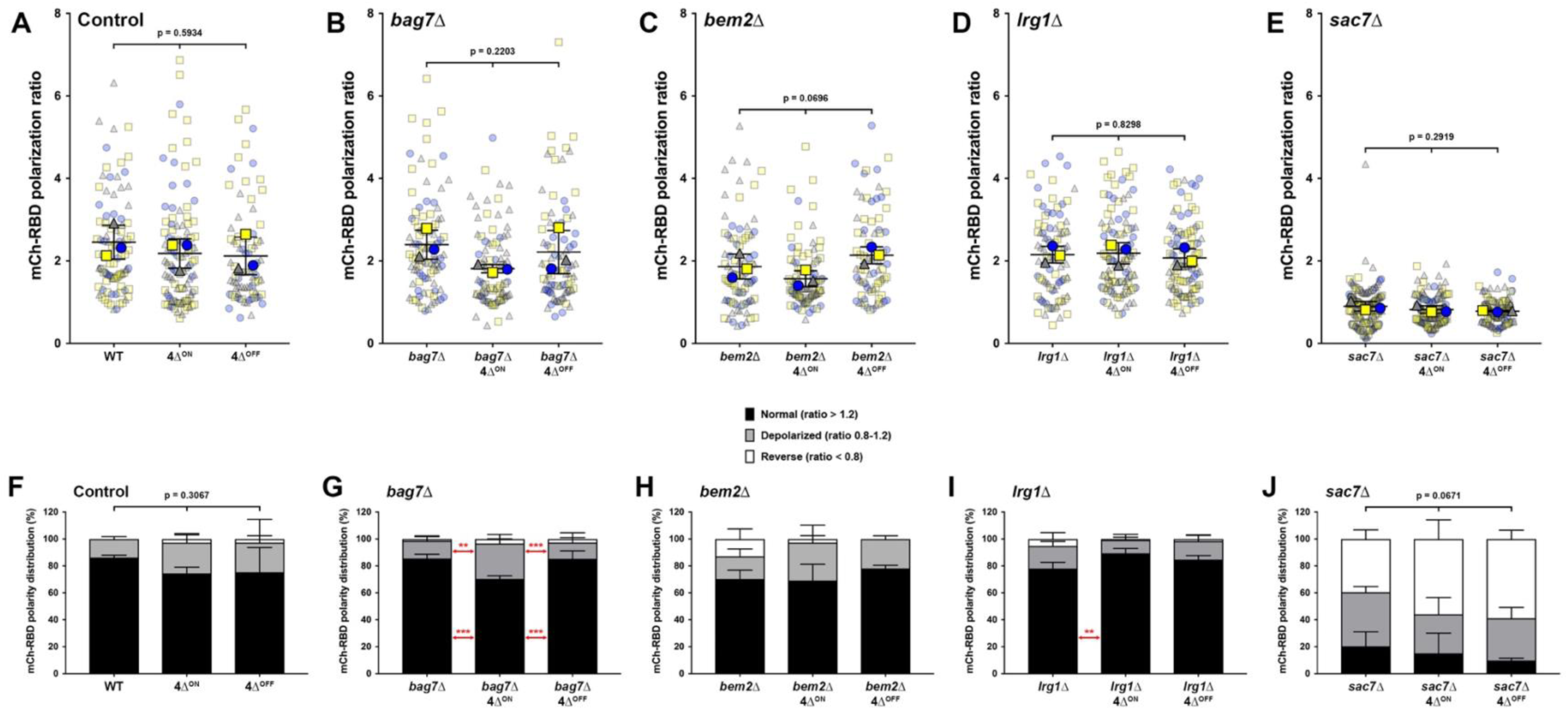
Expanded data from Figure 5 comparing mCherry-Pkc1^RBD^ polarization in WT, 4Δ^ON^, and 4Δ^OFF^ backgrounds for control and Rho1 GAP deletions. (A-E) Polarization ratios for control, 4Δ^ON^ and 4Δ^OFF^ strains without deletion of Rho1 GAPs (A) and in *bag7*Δ (B), *bem2*Δ (C), *lrg1*Δ (D) and *sac7*Δ (E) backgrounds were quantified in budding cells as described in Figure 5B. Polarization ratios were calculated for n=3 biological replicates, with 18-56 cells measured per condition for each trial. One-way ANOVA with Tukey’s multiple comparisons test. (F-J) Classification of control, 4Δ^ON^ and 4Δ^OFF^ strains without deletion of Rho1 GAPs (F) and in *bag7*Δ (G), *bem2*Δ (H), *lrg1*Δ (I) and *sac7*Δ (J) backgrounds as cells with normal (ratio > 1.2), depolarized (ratio between 0.8-1.2), or reversed (ratio < 0.8) polarization of mCherry-RBD. Cells from each trial in panels A-E were categorized based on the above ratios and expressed as a percentage (n=3). Two-way ANOVA with Tukey’s multiple comparisons test, ** p<0.01; *** p<0.001. Note that the left-most column for all polarization ratio (A-E) and classifications of polarity distribution (F-J) are identical to data in Figures 5B and C, respectively, because quantifications and classifications were based on three independent trials of imaging as shown in Figure 5A, where all Rho1 GAP deletion and adaptor mutation strains were imaged in the same session for each of the three trials.

**Supplementary Figure S5:**
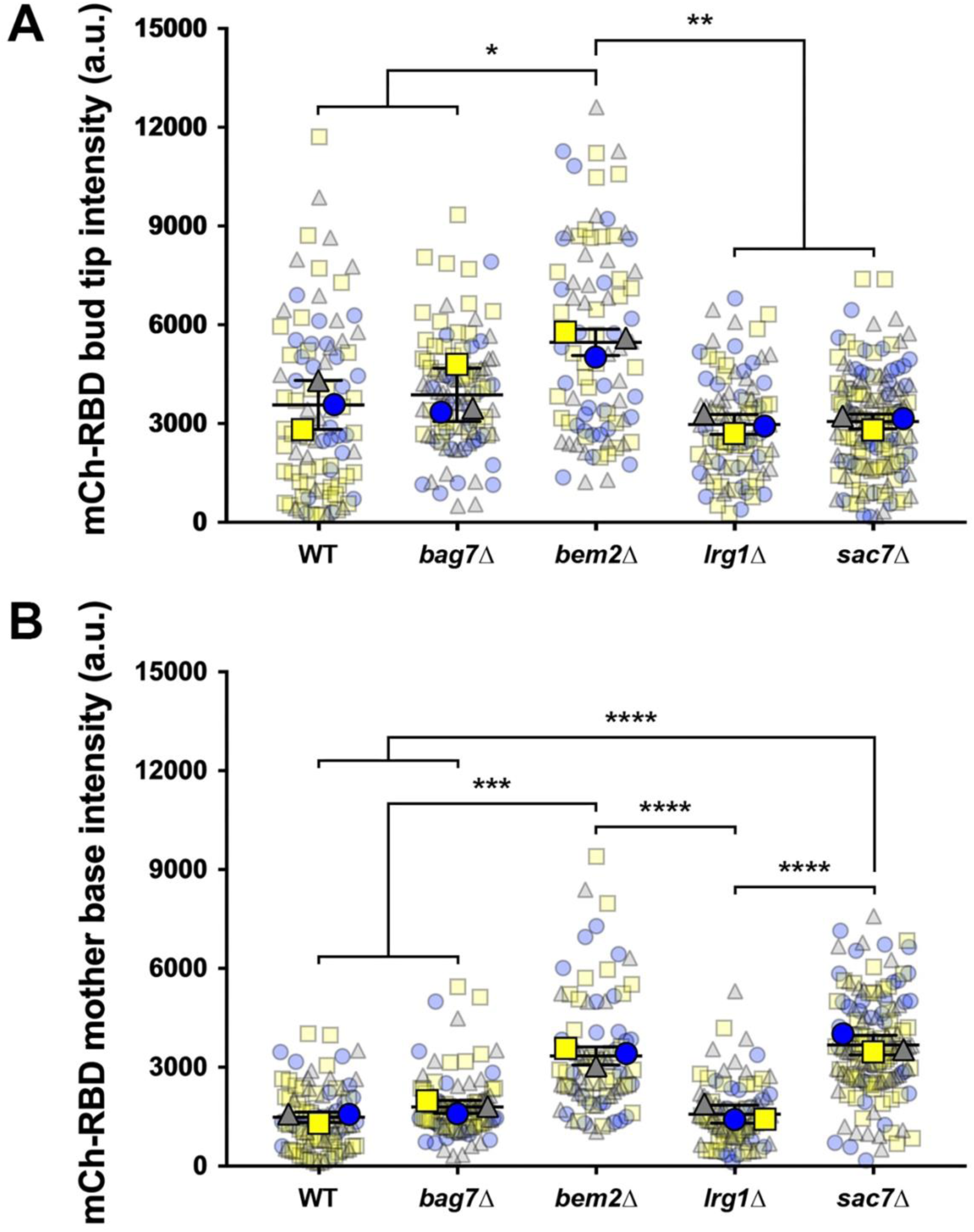
Effect of Rho1 GAP deletion on mother and bud recruitment of mCherry-Pkc1^RBD^. WT, *bag7*Δ, *bem2*Δ, *lrg1*Δ, and *sac7*Δ cells imaged in Fig. 5 for assessment of mCherry-Pkc1^RBD^ polarization were additionally analyzed for peak fluorescence intensity (in arbitrary units, a.u.) at the bud tip (A) and base of the mother cell (B). Fluorescence intensity values were measured for n=3 biological replicates, with 23-56 cells measured per condition for each trial. One-way ANOVA with Tukey’s multiple comparisons test, * p<0.05; ** p<0.01; *** p<0.001; **** p<0.0001.

**Movie 1: Timelapse imaging of Bni1-3GFP in WT cells.** WT cells expressing genetically-encoded Bni1-3GFP as shown in Fig. 6A were imaged by near-TIRF microscopy at a rate of 20 fps. Movie playback is in real-time (20 fps) to show dynamics of cytoplasmic and cortical Bni1-3GFP speckles.

**Movie 2: Timelapse imaging of Bni1-3GFP in *bag7*Δ cells.** *bag7*Δ cells expressing genetically-encoded Bni1-3GFP as shown in Fig. 6A were imaged by near-TIRF microscopy at a rate of 20 fps. Movie playback is in real-time (20 fps) to show dynamics of cytoplasmic and cortical Bni1-3GFP speckles.

**Movie 3: Timelapse imaging of Bni1-3GFP in *bem2*Δ cells.** *bem2*Δ cells expressing genetically-encoded Bni1-3GFP as shown in Fig. 6A were imaged by near-TIRF microscopy at a rate of 20 fps. Movie playback is in real-time (20 fps) to show dynamics of cytoplasmic and cortical Bni1-3GFP speckles.

**Movie 4: Timelapse imaging of Bni1-3GFP in *lrg1*Δ cells.** *lrg1*Δ cells expressing genetically-encoded Bni1-3GFP as shown in Fig. 6A were imaged by near-TIRF microscopy at a rate of 20 fps. Movie playback is in real-time (20 fps) to show dynamics of cytoplasmic and cortical Bni1-3GFP speckles.

**Movie 5: Timelapse imaging of Bni1-3GFP in *sac7*Δ cells.** *sac7*Δ cells expressing genetically-encoded Bni1-3GFP as shown in Fig. 6A were imaged by near-TIRF microscopy at a rate of 20 fps. Movie playback is in real-time (20 fps) to show dynamics of cytoplasmic and cortical Bni1-3GFP speckles.

**Supplementary Table 1:**
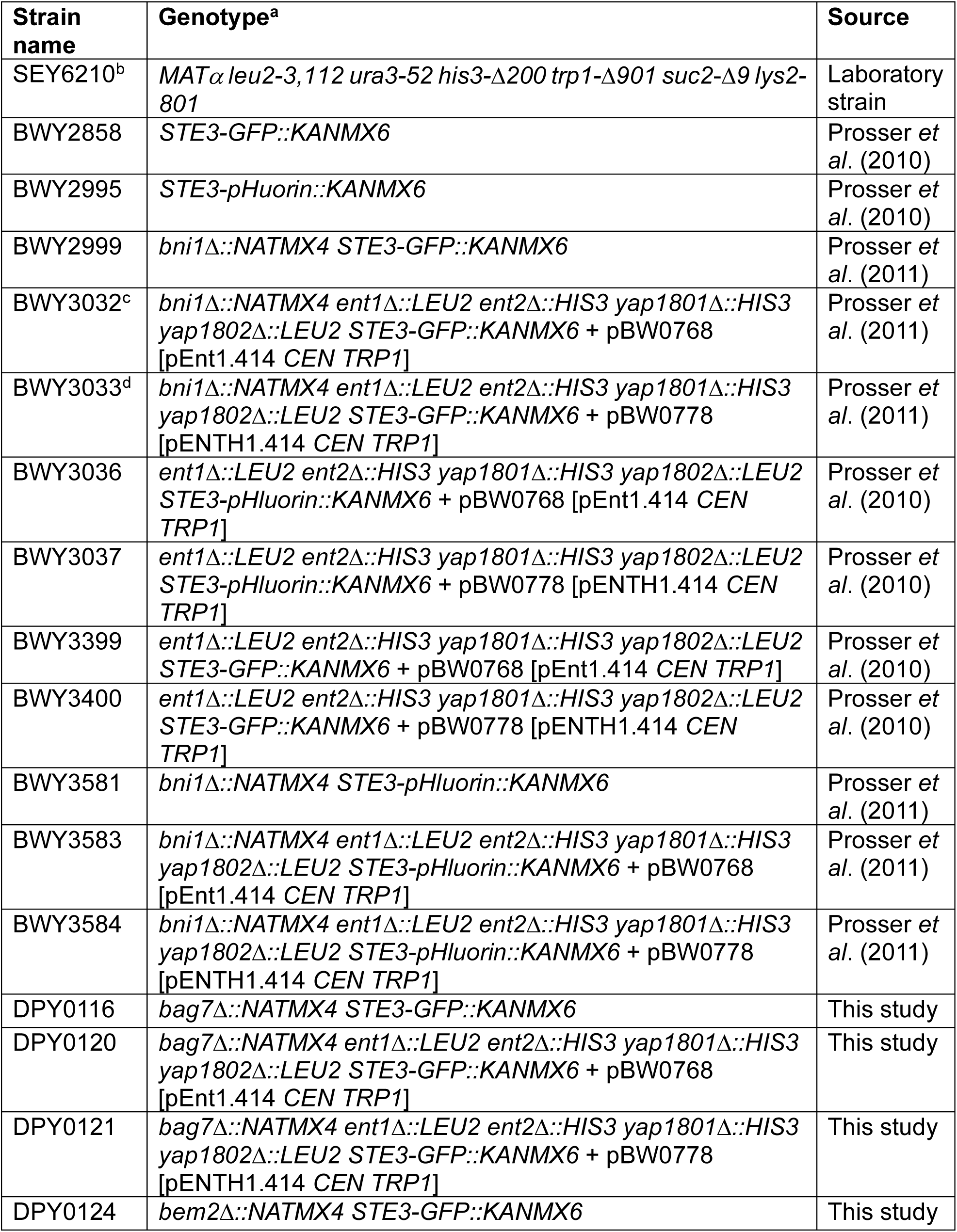
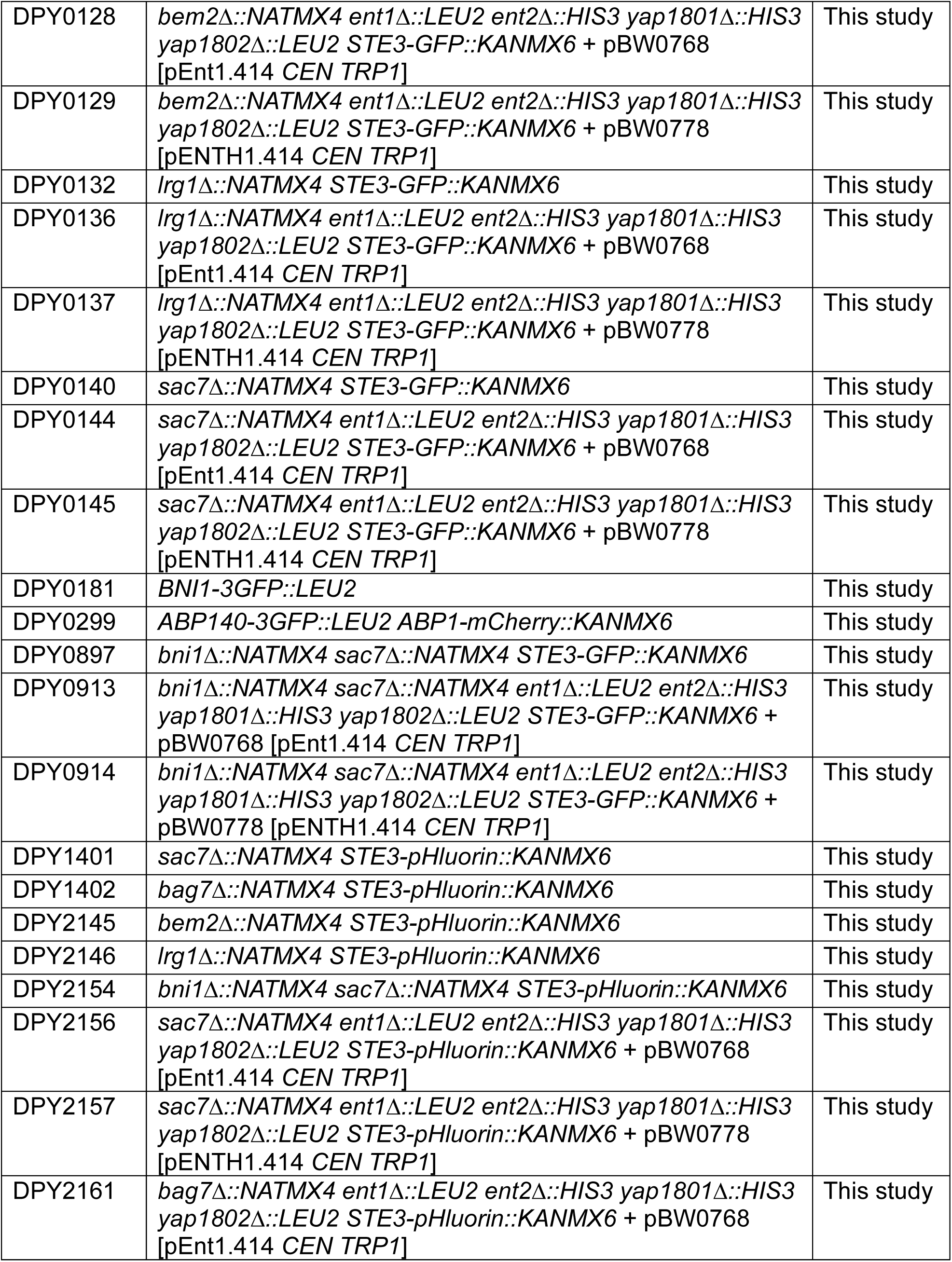
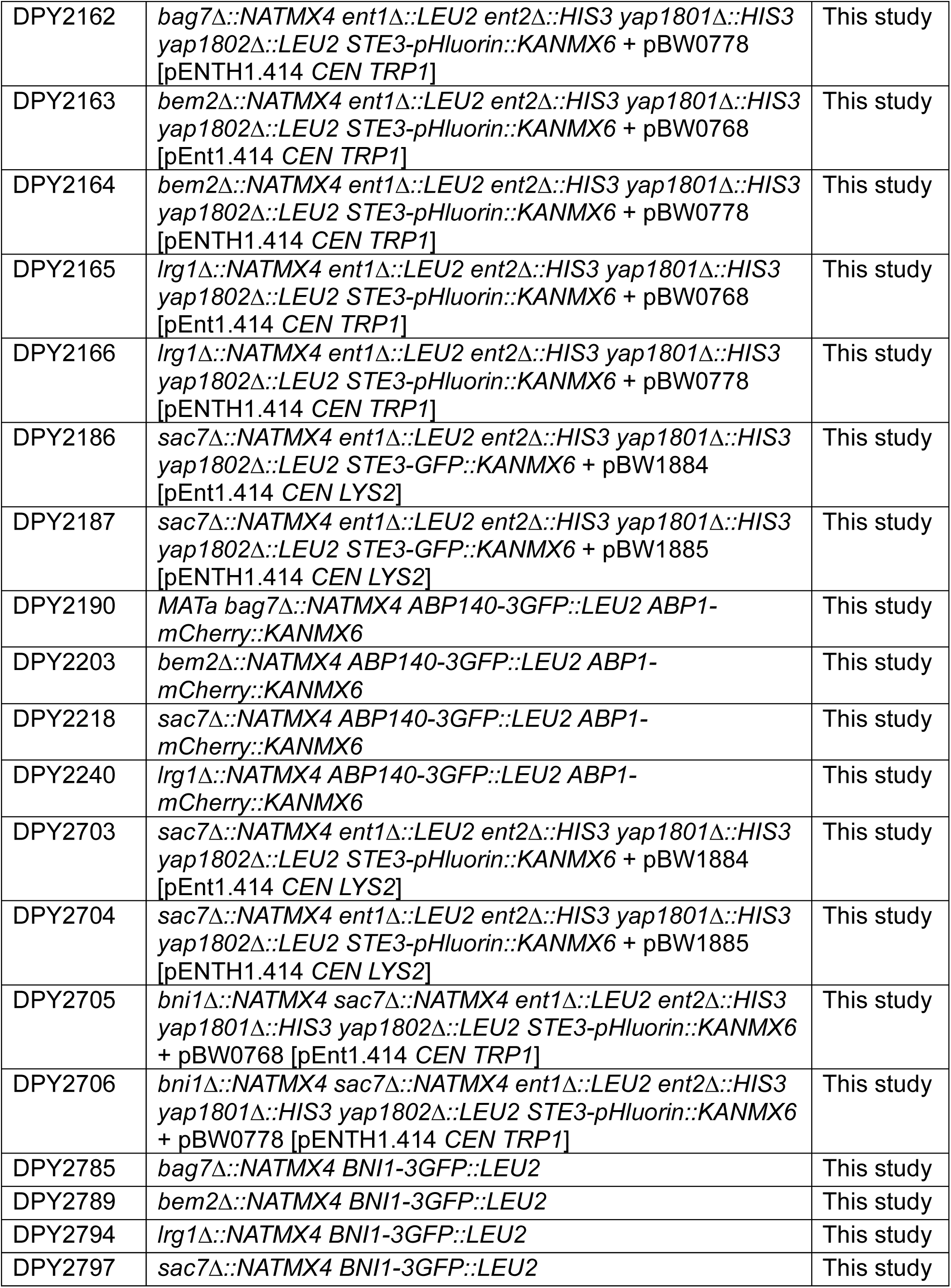
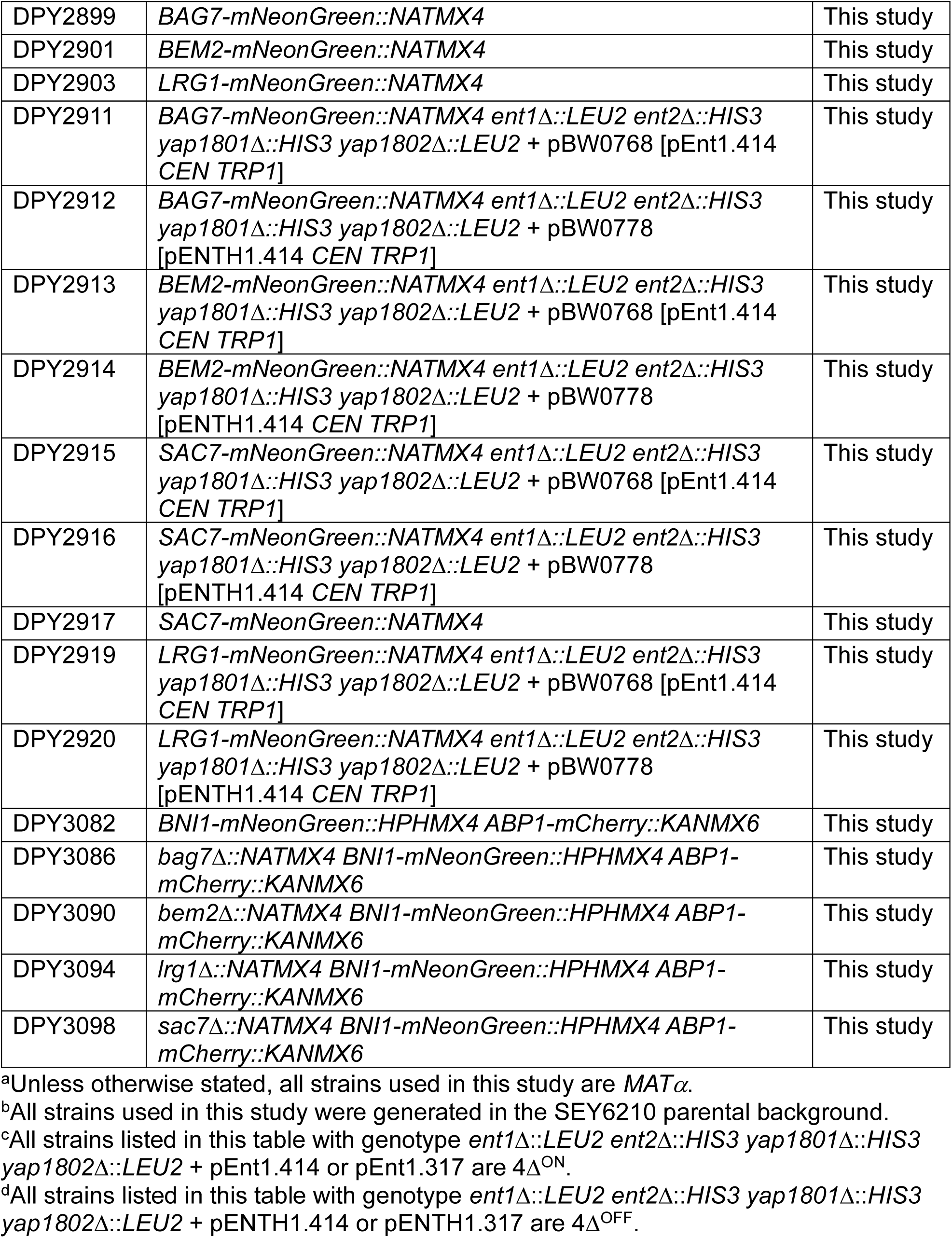
Yeast strains used in this study.

**Supplementary Table 2:** Plasmids used in this study.

| Plasmid name | Details | Description | Source |
| --- | --- | --- | --- |
| pRS317 | empty vector [*CEN LYS2*] | pRS317 | Christianson *et al.* (1992) |
| pRS414 | empty vector [*CEN TRP1*] | pRS414 | Christianson *et al.* (1992) |
| pRS426 | empty vector [2μ *URA3*] | pRS426 | Christianson *et al.* (1992) |
| pB1993 | pRS305::*BNI1*-3GFP [*LEU2*] | pBni1-3GFP [*LEU2*] | Buttery *et al.* (2007) |
| pB1994 | pRS305::*ABP140*-3GFP [*LEU2*] | pAbp140-3GFP [*LEU2*] | Buttery *et al.* (2007) |
| pBW0768 | pRS414::*ENT1* [*CEN TRP1*] | pEnt1 [*CEN TRP1*] | Maldonado-Báez *et al.* (2008) |
| pBW0778 | pRS414::*ent1*(aa1-151, ENTH domain only) [*CEN TRP1*] | pENTH1 [*CEN TRP1*] | Maldonado-Báez *et al.* (2008) |
| pBW1884 | pRS414::*ENT1* [*CEN LYS2*] | pEnt1 [*CEN LYS2*] | Prosser *et al.* (2011) |
| pBW1885 | pRS414::*ent1*(aa1-151, ENTH domain only) [*CEN LYS2*] | pENTH1 [*CEN LYS2*] | Prosser *et al.* (2011) |
| pBW2053 | YE<sub>p24</sub>::ROM1 [2μ *URA3*] | pRom1 [2μ *URA3*] | Prosser *et al.* (2011) |
| pDP0480 | pRS317::*GFP-RHO1* [*CEN LYS2*] | pGFP-Rho1 [*CEN LYS2*] | This study |
| pDP0481 | pRS317::*GFP-RHO1*<sup>Q68H</sup> [*CEN LYS2*] | pGFP-Rho1<sup>Q68H</sup> [*CEN LYS2*] | This study |
| pDP0551 | pRS317::*P<sub>MET25</sub>-mCherry-Pkc1<sup>384-632</sup>-T<sub>CYC1</sub>* [*CEN LYS2*] | pmCherry-Pkc1<sup>RBD</sup> [*CEN LYS2*] | This study |
| pDP0764 | pRS414::*SAC7* [*CEN TRP1*] | pSac7 [*CEN TRP1*] | This study |
| pDP0765 | pRS414::*sac7*<sup>R173K</sup> [*CEN TRP1*] | pSac7<sup>R173K</sup> [*CEN TRP1*] | This study |
| SP301 | GFP-Rho1 [*CEN URA3*] | pGFP-Rho1 [*CEN URA3*] | Yoshida *et al.* (2009) |
| SP307 | GFP-Rho1<sup>Q68H</sup> [*CEN URA3*] | pGFP-Rho1<sup>Q68H</sup> [*CEN URA3*] | Yoshida *et al.* (2009) |

